# Soil Resistomes in a Tropical Watershed are Indirectly Structured by Bacterial Community Interactions with Soil Properties

**DOI:** 10.64898/2026.06.18.733189

**Authors:** Wesley J. Sparagon, Sean Lary, Makana T. Ioh, Andrew Lin, Ishwora Dhungana, Christian R. Fullmer, Christy R. Handel, Roshan Paudel, Juli Burden, Justene N. Deubel, Malissa Ann G. Tayo, Francisca E. Rodriguez, Sean O.I. Swift, Kirsten K. Nakayama, Tai M. Maaz, Nhu H. Nguyen

## Abstract

Soils are recognized as reservoirs of antibiotic resistance genes (ARGs) with the potential to transfer to clinical pathogens, creating antimicrobial resistance (AMR) that poses a threat to human health. While large-scale AMR surveys have profiled how diverse biomes shape soil resistomes, less is known about the influence of specific soil properties. Here, we combined metagenomics and 16S rRNA amplicon sequencing with isolate-based approaches to investigate drivers of soil AMR across a tropical watershed from beach to mountaintop in Waimea Valley, O’ahu, Hawai□i. We leveraged functional- and taxonomic-classification of resistances to unravel how soil properties interact with bacterial taxa to structure resistomes. Metagenomic- and isolate-resistomes showed remarkable consistency, including a general gradient of increasing AMR from ridge to beach. Resistome functional composition was significantly correlated with total bacterial community structure. The relationship between resistances and soil properties was primarily dictated by taxonomic composition of each resistance. Rifampin- and Vancomycin-ARGs associated with Actinomycetes negatively correlated with soil physical properties, while resistant genes and isolates from Gammaproteobacteria positively correlated with enzymatic activity metrics. These findings indicate that soil properties structure the resistome indirectly through taxonomic filtering of microbial hosts and challenge the notion that AMR is decoupled from phylogenetic relatedness.

## INTRODUCTION

Antimicrobial resistance (AMR) is widely recognized as a global threat to human health and safety. Since the advent of antibiotics, the pervasiveness of AMR among microorganisms has increased due to selective pressures from antibiotic use in healthcare and agriculture. Globally, AMR has resulted in 700,000 deaths per year[1], prompting organizations such as the World Health Organization to deem it a large-scale, immediate concern for human health and safety[2].

Soils are reservoirs of clinically relevant microbes and antimicrobial resistance genes (ARGs)[3]. Antibiotic production by soil microorganisms imposes strong selective pressures, which may be amplified by anthropogenic inputs of clinical antibiotics, collectively driving the emergence and persistence of antibiotic resistance[4, 5]. Because soil-borne ARGs can be horizontally transferred to clinical pathogens, soil resistomes (the resistances harbored, transferred, and maintained among microorganisms) represent a potential source of AMR infections and could contribute to human mortality worldwide[3, 6, 7]. However, the ecological and environmental drivers shaping soil resistome composition and abundance remain poorly understood.

Soil properties govern the assembly and structure of the soil microbiome[8], yet much less is known about their influence on the soil resistome. Global surveys of soil resistomes indicate temperature and pH as key mediators of resistome composition[9]. Across forest biomes, soil pH, soil moisture, and mean annual temperature have been identified as key mediators of the soil resistome[10]. In manure fertilized soils, soil pH (acidic, near-neutral, and alkaline) was shown to exert the most influence over resistomes[11]. Elevated metal content has also been shown to enrich AMR via coselection of ARGs and enhanced horizontal gene transfer of ARGs between organisms[12]. Beyond soil properties, biotic factors such as above-ground herbaceous communities[10] and bacterial community taxonomic composition[13] have also been shown to significantly structure soil resistomes. These findings suggest that a similar set of principal variables govern the assembly of both the soil microbiomes as well as the resistomes harbored within them.

Global resistome surveys have revealed broad patterns in antimicrobial resistance and the factors that contribute to these resistances[14], but their large scale and methodological limitations restrict more specific insights into soil-associated drivers of AMR. These patterns might be masked across global trends due to a lack of replication in specific environments and the large differences between highly divergent biomes. Furthermore, environmental AMR surveys often utilize either culture-dependent or culture-independent methods, with few studies integrating both. It is remiss to scrutinize bacterial soil populations through solely culture-dependent or -independent lenses, as they are often complementary[15, 16]. AMR within microbial communities are complex processes which require multiple approaches. Therefore, a comprehensive synthesis that integrates both biotic factors and abiotic soil properties is necessary to better understand the persistence of antimicrobial resistance in soils.

To address these knowledge gaps, we employed a two-pronged approach combining culture-independent molecular and culture-dependent methods to survey AMR across the Waimea tropical watershed on Ocahu, Hawaici. This watershed begins at a remote, high-elevation ridge, descends through Waimea Valley and the Waimea Botanical Garden, and ultimately flows into coastal rivers and an estuarine system before transecting Waimea Bay[17, 18]. Waimea Valley serves as a valuable natural model for examining the potential drivers of the soil resistome. This watershed exhibits strong gradients not only in elevation and rainfall, but also substantial changes in soil order and mineralogy, across a relatively small geographic area of a tropical moist to wet forest. This allows us to control for the strong influence of biome-type on the soil resistome and directly compare how soil properties might influence the soil resistome. This study builds the connections between resistome structure and soil properties using culture-independent metagenomic and amplicon datasets, combined with culture-dependent isolates, across seven sites along the acclivity of the Waimea watershed.

## MATERIALS AND METHODS

### Site description and sampling

We sampled soils across the Waimea Valley model tropical watershed ecosystem on the Island of Ocahu, Hawaici[17, 19] with modest gain in elevation (0–682 m), a strong precipitation gradient from beach (1.1 m rainfall/y) to ridge (4.6 m rainfall/y), and varying soil characteristics (Table S1). The watershed is subjected to human exposure of varying degrees: the lower elevation portion includes a public beach park, botanical garden, and waterfall that are frequented by tourists. Human traffic dissipates dramatically past the waterfall between the mid-valley and ridge site. Sampling sites along the transect are as follows: Ridge, Midvalley, Waterfall, Confluence, Entrance, Estuary, Beach (Figure S1).

Four surface soil cores (5 × 5 cm) were taken at each sampling site and homogenized into a single sample. This approach was repeated three times (three replicates), 5 m apart at each site. A total of 21 samples were taken across the transect. Samples were kept cool and frozen at −20 °C until further processing. The soils were subset into three portions to measure antibiotic resistance of resident microbes, bacterial community structure, and soil characteristics. Soil characteristics for each sample were measured at the Soils & Ecosystems Lab at the University of Hawai□i at Ma□noa (UH Ma□noa). The Hawai□i Soil Health test[20, 21] included physical (aggregate stability, water holding capacity, moisture), chemical (pH, P, K, Ca, Mg, Na, total C, total N, high molecular-weight C, dissolved organic C, potentially mineralizable N, dissolved organic N) and biological (CO_2_ burst, β-glucosidase, β-glucosaminidase) properties. We also measured additional trace elements and heavy metals associated with these soils (Pb, V, Cr, Fe, Zn, Cu, Mn, Co, and Ni) on a Perkin-Elmer ICP-OES using nitric acid extractions (Table S2).

### Bacteria isolation and antibiotic susceptibility test

Resistant soil bacteria were isolated from each sample on selective Luria-Bertani (LB) agar, supplemented with one of nine antibiotics, each belonging to a different class: ampicillin (Penicillins, 50 µg/mL), cefepime (Cephaloporins, 2 µg/mL), erythromycin (Macrolides, 1 µg/mL), kanamycin (Aminoglycosides, 50 µg/mL), levofloxacin (Fluoroquinolones, 3 µg/mL), meropenem (Carbapenems, 0.25 µg/mL), rifamycin (Rifampicins, 2.5 µg/mL), tetracycline (Tetracyclines, 50 µg/mL), and vancomycin (Glycopeptides, 4 µg/mL). Isolates were streaked onto LB plates to obtain 865 cultures and maintained on 20% glycerol stock at −80 °C. To confirm and standardize antibiotic susceptibility, each isolate was tested using the Kirby-Bauer disk assay[22]. Isolation and assay details are presented in the Supplementary Methods and Table S3.

### DNA extraction, amplicon library preparation, and sequencing

DNA was extracted from 0.25 g of homogenized soil using the DNEasy PowerSoil Kit (Qiagen, USA) following manufacturer’s instructions. A 16S rRNA gene amplicon library for bacterial community profiling was prepared following a two-step procedure[23]. The first step amplifies the V4 region of the bacterial and archaeal 16S rRNA gene using the primers 515F and 806R[24, 25]. Amplified products were cleaned using solid-phase reversible immobilization (SPRI) magnetic beads, and the cleaned products were used for the second PCR to attach an 8 bp barcode and Illumina sequencing adapters to each end of the amplicon. PCR details are provided in Supplementary Methods. The barcoded products were cleaned using SPRI magnetic beads as above, quantified using the Qubit 3.0 fluorometer (Thermo Fisher Scientific, USA), combined in equimolar concentration. A PCR negative control and a mock community as positive control were added along with the experimental samples according to the guidelines in Nguyen et al., 2015[26]. The final library was sequenced using Illumina MiSeq 300PE at the Advanced Studies in Genomics, Proteomics and Bioinformatics core facility at the University of

Hawai□i at Ma□noa. DNA extracts from the same samples were prepared for metagenomic shotgun sequencing with Illumina DNA Prep Kit with Nextera XT v2 Index Kit Set A and sequenced using Illumina NextSeq 500 (2 × 150bp) at the University of Hawai□ i at Ma□noa Genomics and Bioinformatics Laboratory. The run yielded 2,362,961–10,355,608 reads per sample.

Isolates were identified using high-throughput sequencing[27]. DNA was extracted from each isolate and PCR was performed using a two-step, dual barcoding approach as above using the crude extract as the template, with MyTaq Red mastermix (Meridian Bioscience, USA), and the primer pairs 515F[25] and 926R[28] that amplify the V4-V5 region. DNA extraction and PCR details provided in Supplementary Methods. Barcoded products from all isolates were pooled at equimolar concentration, cleaned using SPRI beads to remove excess primers, and sequenced on the Illumina MiSeq 300PE as above.

### Bioinformatics

Amplicon sequence data processing followed standard protocol for sequence quality control using QIIME2 v2021.2[29]. Pipeline details are provided in Supplementary Methods. Sequences classified as “unassigned”, “mitochondria”, “chloroplast” and “Archaea” were removed since they were not of primary interest in this study. One amplicon sequence variant (ASV), a *Bosea* sp., showed up as highly abundant in the negative control and thus was removed from the whole dataset. Any ASV that contained less than 10 sequences total were also removed to minimize noise[26]. All samples were then rarefied to 85,292 sequences, and the final ASV contingency table was exported for analyses in R.

A similar bioinformatic workflow with additional steps was used to process sequences from isolates. First, the sequences were processed as above to obtain high-quality ASVs. A large proportion of the pure isolates, especially of certain taxa (e.g. *Microbacterium, Paenarthrobacter, Flavobacterium, Chryseobacterium, Pseudomonas*), produced multiple ASVs per isolate that had similarities between 98-99% to each other. We think that this is due to the polymorphic nature of these taxa at the V4-5 region[30]. Given this, we chose to cluster this dataset at 99% similarity to reduce the number of OTUs that might appear in a sample.

Sequences classified as “unassigned”, “mitochondria”, “chloroplast”, “*Archaea*”, and the unculturable taxa “*Planctomycetota*”, or “*Acidobacteria*” were removed. Further details are provided in the Supplementary Methods. After removal of isolates not meeting purity criteria, along with those that were not sequenced successfully, our final dataset contained 771 pure isolates with identification (Table S4).

Metagenomic reads were processed using the AMR++ v3.0 pipeline[31]. In brief, raw data was filtered using trimmomatic[32], deduped, and aligned to the MEGARES[33] database using BWA[34]. Reads were taxonomically classified using Kraken2[35] with the “standard nt” refseq database. MEGs requiring single nucleotide polymorphism (SNP) confirmation were further processed using the AMR++ v3.0 built-in SNP Confirmation workflow. MEGs represented in only one sample were removed and output MEG counts were normalized to sequencing depth by calculating the MEG sequences/million raw reads value, resulting in a final gene abundance table containing 50 identified MEGs (Table S5). For consistency with the literature, all MEGs will be described as “ARGs” throughout the results and discussion. Detailed AMR++ workflow specifics can be found in the associated github repository (https://github.com/nnguyenlab/waimea-watershed-resistome).

### Statistical analyses

Statistical analyses were performed in R v4.4.2[36]. Weighted UniFrac distance between microbial communities (16S rRNA gene amplicons) was extracted from ‘qiime2R v0.99.6’[29]. The normalized (reads per million) metagenomic MEG abundance table was used to generate a Bray-Curtis dissimilarity matrix for downstream multivariate analysis. Isolate resistance data was used to generate a table of the total number of isolates resistant to each of the assayed antibiotics in each sample. This isolate table was then used to generate a Bray-Curtis dissimilarity matrix for downstream multivariate analysis. To evaluate taxonomic composition of resistome datasets, each read mapping to a known ARG (metagenomes) and each isolate exhibiting AMR (cultures) was assigned a taxonomy via kraken2 and QIIME2, respectively. The relative abundance of taxonomic classes in each ARG/isolate and for the total resistome in each sample was used to generate Bray-Curtis dissimilarity matrices for downstream multivariate analysis.

Packages used for downstream analysis included Vegan v2.6-10[37] for beta diversity analysis and ordinations, randomForest v4.7-12 for all random forest (RF) analysis[38], RcmdrMisc v2.9-for spearmen correlation analysis[39], and ComplexHeatmap v2.22.0 for correlation visualizations[40]. All data was assessed for normality prior to parametric statistical testing and converted to a normal distribution via the log_10_ transformation whenever indicated. Non-normal data were tested using non-parametric Kruskal-Wallis tests and random forest analysis. P-value multiple-comparison adjustment was performed using the Benjamini-Hochberg correction whenever indicated[41].

## RESULTS

### Resistome Functional and Taxonomic Composition

Soil resistome richness from *in situ* metagenomes and *in vitro* cultured isolates exhibited consistent patterns across sample sites. Richness was similar across sites for the total bacterial community (16S rRNA gene amplicons, Kruskal-Wallis, p>0.05, Figure S2A), metagenomic resistomes (ARG abundances, Kruskal-Wallis, p>0.05, Figure S2B), and isolate resistomes (resistant isolate counts, ANOVA, p>0.05, Figure S2C). Resistant isolate counts were significantly positively correlated with ARG abundances (Spearman’s Rank Correlation, Spearman’s ρ=0.5052, p=0.039, Figure S2D). *In situ* metagenomes had 50 unique ARGs spanning 16 functional classes (Table S5). Dominant ARG classes included Rifampin-resistance (354 reads per million total reads), Macrolides, lincosamides, streptogramines (MLS)-resistance (151 reads per million), Aminoglycoside-resistance (57 reads per million), and Glycopeptide-resistance (49 reads per million) (Figure S3A). Isolate resistomes were dominated by Ampicillin-resistance (32 isolates), Vancomycin (Glycopeptide)-resistance (19 isolates), and Erythromycin (MLS)-resistance (17 isolates) (Figure S3B). The most prevalent resistant bacteria taxa were *Pseudomonadales* (28.7%) and *Micrococcales* (27.7%), followed by *Flavobacteriales* (10.1%), Bacillales (7.4%), Burkholderiales (5.7%), Enterobacterales (4.2%), Caulobacterales (3.9%), *Xanthomonadales* (3.9%), *Rhizobiales* (2.8%), *Sphingobacterales* (2%), and *Corynebacteriales* (1.9%) (Figure S4).

Multivariate analysis indicated that bacterial community structure is linked to resistome structure across the sites. The functional and taxonomic classification of ARGs/isolates, along with taxonomic profiling of total bacterial communities, yielded three groups of data analyzed in tandem: 1) “resistome functional composition”, reflecting the abundance of resistance types (ARGs/isolates) in each sample (Figure S3), 2) “resistome taxonomic composition”, reflecting taxonomic assignment of all resistant reads/isolates in a sample, and 3) “total bacterial taxonomic composition”, reflecting the relative abundance of bacterial taxa in each soil sample derived from 16S rRNA gene amplicon sequencing of the entire bacterial community. Site had a significant effect on resistome taxonomic composition in the metagenomes (PERMANOVA, p=0.005, R^2^=0.56318, F=3.0083, Figure S5A) and isolates (PERMANOVA, p=0.033, R^2^=0.39234, F=1.5065, Figure S5B). Site also explained similar or more variation in resistome taxonomic composition compared to resistome functional composition (Figure 1A and 1C, Figure S5C and S5D). Additionally, PERMANOVA R^2^ values indicated that total bacterial taxonomic composition correlated more strongly with site than resistome taxonomic compositions (PERMANOVA, p=0.001, R^2^=0.69761, F=5.3829, Figure 1A and 1B). This suggested that site properties had the strongest influence on total bacterial taxonomic composition, which in turn influenced downstream resistome taxonomic composition and functional composition. Total bacterial taxonomic composition correlated significantly with resistome functional compositions in both the metagenomes (Mantel Test, r=0.19, p=0.047, Figure 1B) and isolates (Mantel Test, r=0.275, p=0.036, Figure 1D), further supporting this concept. Together, these lines of evidence indicate that resistome structure in both datasets is driven by variations in the soil bacterial community, which are in turn likely driven by changing soil properties across the transect.

**Figure 1:**
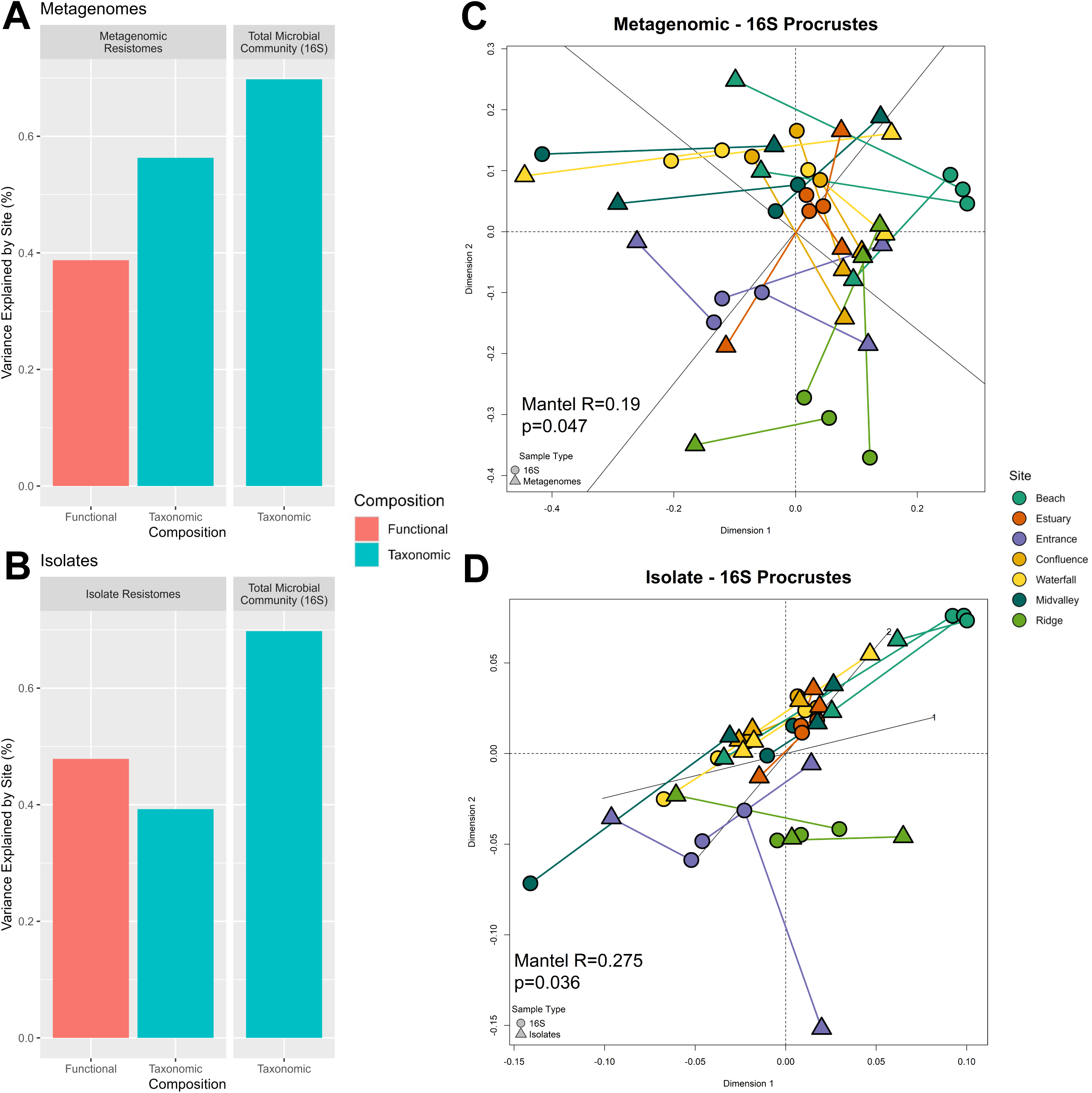
Multivariate patterns in the soil resistome. Barplots depict the variance explained by Site (derived from PERMANOVA R^2^ values) when tested against the functional- and taxonomic-composition of the resistomes and total bacterial taxonomic composition, in the **A)** metagenomes and **C)** isolates. Significant multivariate correlations (Mantel test, p<0.05) between total bacterial taxonomic composition and both **B)** metagenomic resistome functional composition and **D)** isolate resistome functional composition are depicted as procrustes plots. Lines connect paired samples, and point shape corresponds to sample type.

### Soil Properties that Structure the Soil Resistome

To identify how soil properties may be connected to soil resistome structure, we performed Spearman’s Rank Correlation between ARG abundances and soil properties (Figure 2), and resistant isolate counts and soil properties (Figure 3). Spearman’s rank correlation coefficient values were visualized in biclustering heatmaps to identify consistent AMR-soil property relationships. The resulting biclustering heatmaps grouped both metagenomic resistances and isolate resistances into three clusters via k-means clustering. Resistances within these clusters exhibited consistent correlations with specific groups of soil properties. We will hereafter refer to these clusters as “ARG Cluster 1-3” and “Isolate Cluster 1-3”. The soil properties formed five clusters via k-means clustering, hereafter referred to as “groups”. Soil properties formed similar groups across the two independent datasets: the “Enzymatic Activity” group was connected to microbial metabolism such as B-glucosidase and B-glucosaminidase activities, dissolved organic carbon:dissolved organic nitrogen (DOC:DON) ratios, and macronutrients such as P and K; the “Physical Properties” group (also as a subgroup in the isolate dataset) included aggregate stability, water holding capacity, and gravimetric water content; and the “Trace Elements” group which contained trace elements and metals such as Pb, V, Cr, Fe, Zn, Cu, Mn, Co, and Ni.

**Figure 2:**
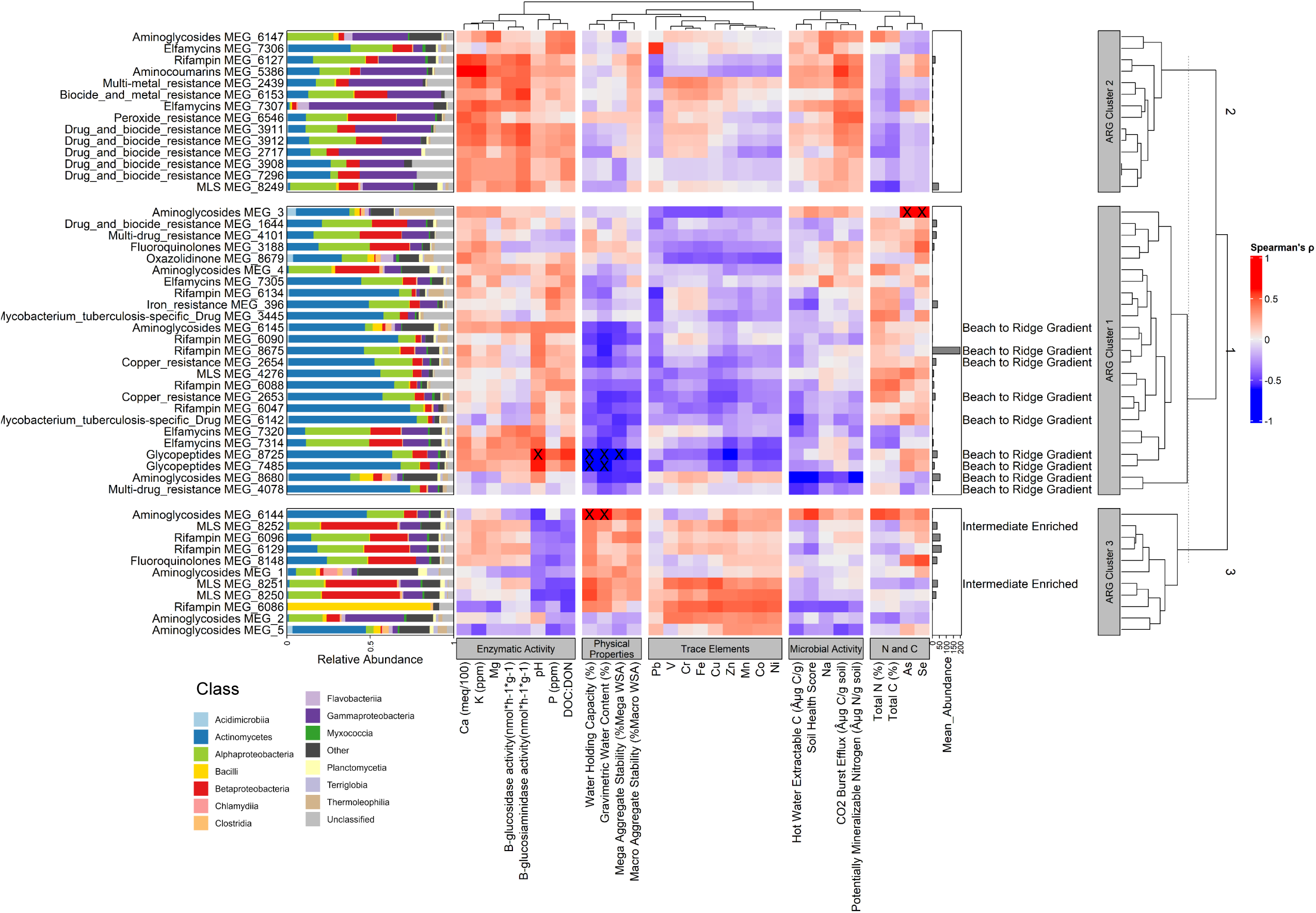
Soil properties that structure metagenomic resistome. A biclustering heatmap of Spearman’s Rank Correlation ρ values between individual ARGs (rows) and soil properties (columns). Significant correlations (Benjamin-Hochberg adjusted p<0.05) are marked with an “X” over cells. Soil properties are split into similarly-behaving clusters labeled at the base of the heatmap and ARGs are split via k-means clustering into similarly-responding clusters labeled “ARG Cluster 1-3” to the right of the heatmap. ARGs are labeled on the left according to their resistance classes and associated accession number. To the right of ARG labels, their taxonomic composition is displayed as relative abundance of bacterial classes. Classes were considered “Other” if the mean relative abundance of said class was < 1%. The mean normalized abundance of each ARG is displayed as a barplot directly to the right of the heatmap. Indicator ARGs are labeled according to their group to the right of these barplots.

**Figure 3:**
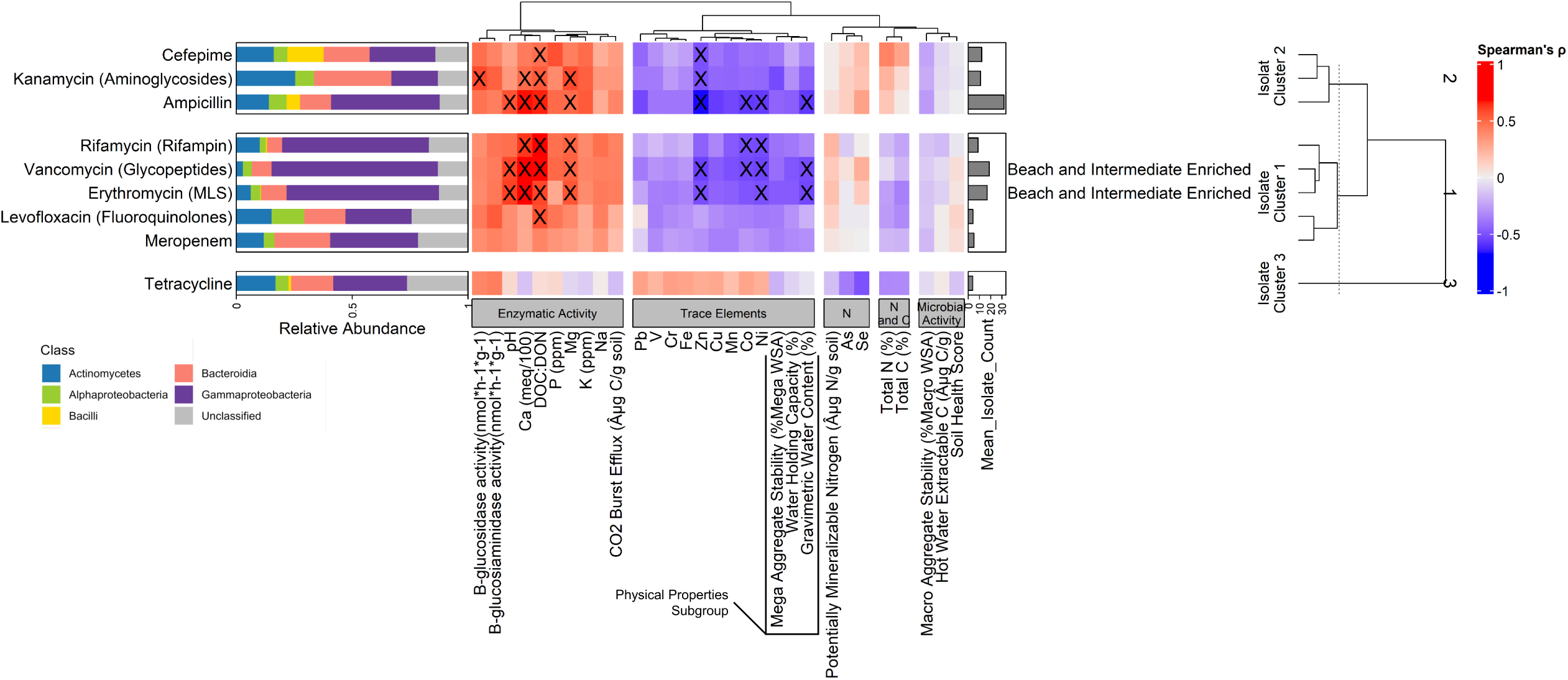
Soil properties that structure isolate resistome. A biclustering heatmap of Spearman’s Rank Correlation ρ values between antibiotic isolate resistance counts (rows) and soil properties (columns). Significant correlations (Benjamin-Hochberg adjusted p<0.05) are marked with an “X” over cells. Soil properties are split into similarly-behaving clusters labeled at the base of the heatmap and isolate resistances are split via k-means clustering into similarly-responding clusters labeled “Isolate Cluster 1-3”. Isolate resistances are labeled on the left according to their antibiotic resistances. To the right of resistance labels, their taxonomic composition is displayed as relative abundance of bacterial classes. The mean resistant isolate count of each antibiotic resistance is displayed as a barplot directly to the right of the heatmap. Isolate resistances that significantly responded to site are labeled according to their group to the right of these barplots.

The three major groups of soil properties, Physical Properties, Trace Elements, and Enzymatic Activities were important factors connected to the structure of the soil resistome. The abundance of metagenomic ARG Cluster 1 negatively correlated with the Physical Properties group (lm, p=0.004, R^2^=0.324, Figure 4A) and showed a significant decreasing beach-to-ridge gradient (ANOVA, p=0.018, F=3.841, Figure 4B). ARG Cluster 1 included the most abundant ARGs in the dataset, namely Rifampin MEG 8675, Aminoglycoside MEG 8680, and Glycopeptide MEG 8725 (Figure 2). This cluster also included all ARGs exhibiting a beach-to-ridge decreasing gradient identified by random forest (RF) analysis (Mean Decrease in Accuracy > 0.006, Figure S6, Figure 5A) such as Aminoglycoside, Copper, Glycopeptide, and Rifampin ARGs. ARG Clusters 2 and 3 showed significant positive correlations with the Enzymatic Activity group (lm, p=0.027, Figure 4C) and Trace Elements group (lm, p=0.021, Figure 4E) across the transect, respectively. This relationship resulted in higher abundance at intermediate sites in the transect (Figure 4D and 4F) and was corroborated by random forest which identified intermediate enriched MLS-resistant genes in ARG Cluster 3 (MLS MEG 8251 and 8252, Mean Decrease in Accuracy > 0.006, Figure S6, Figure 5B).

**Figure 4:**
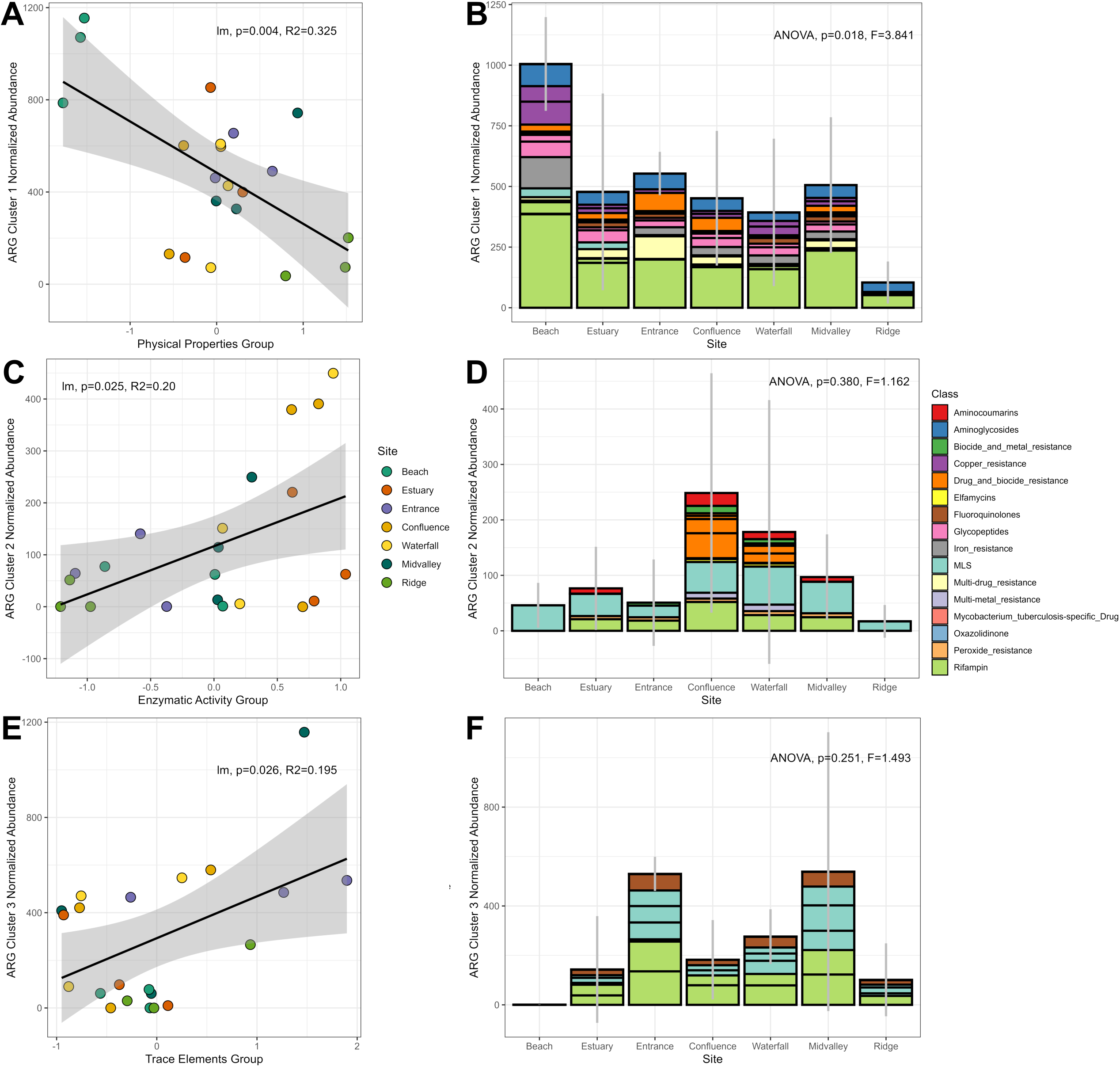
Significant correlations between soil properties and ARG Clusters yield emergent patterns in the *in situ* soil resistome across sites. Significant correlations (lm, p<0.05) between ARG Cluster abundances and soil property groups defined in Figure 2 are displayed in the left column for **A)** Cluster 2, **C)** Cluster 1, and **E)** Cluster 3. Indices for the soil groups were calculated by summing the z-score normalized values for every variable in a group. Mean normalized abundances of ARG Clusters across sites are displayed in the righthand column for **B)** Cluster 2, **D)** Cluster 1, and **F)** Cluster 3. Error bars indicate ± standard deviation, and bar colors indicate the ARG resistance classes in each cluster.

**Figure 5:**
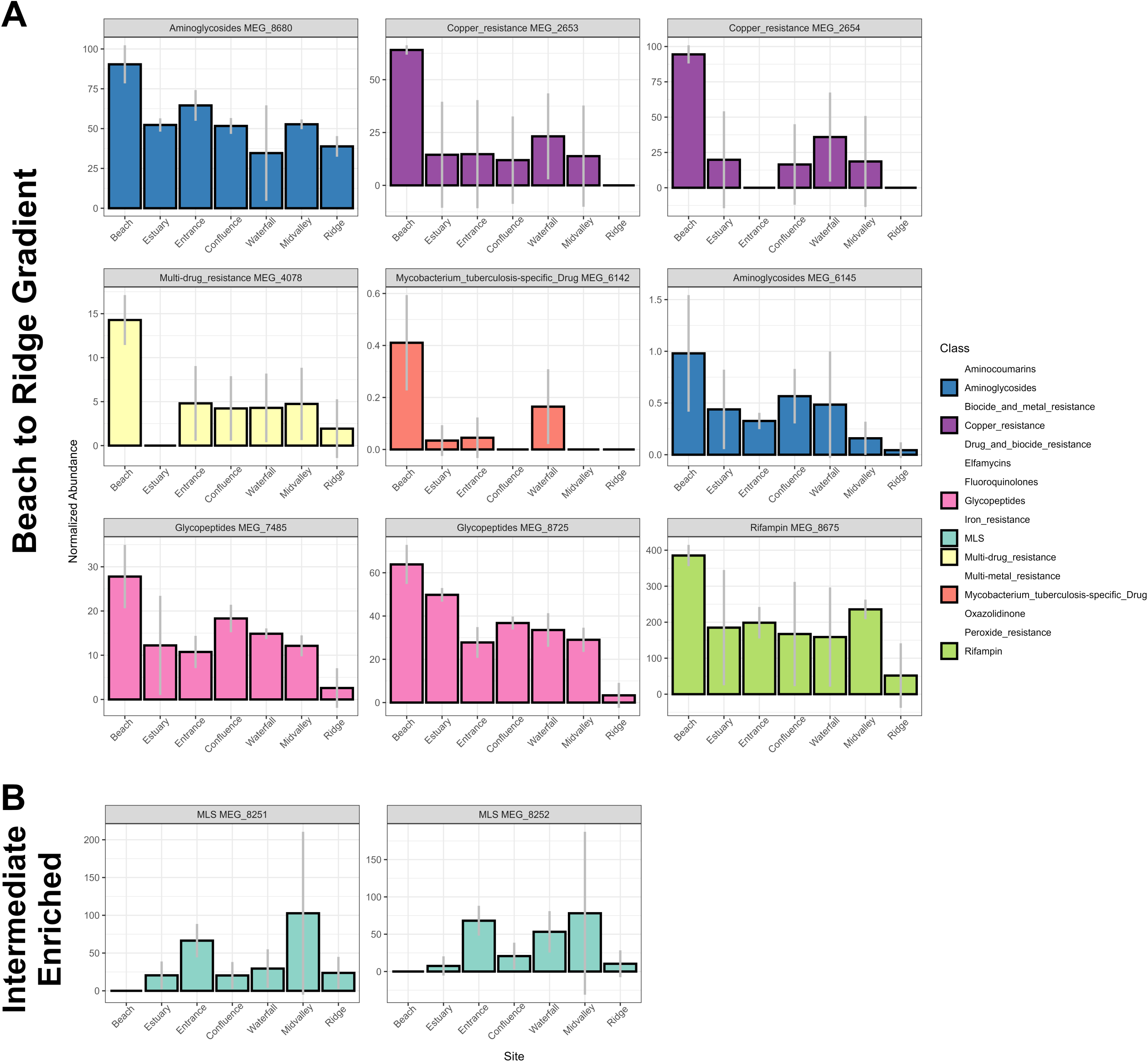
Site indicator ARGs identified via random forest machine learning. Bar heights represent mean normalized abundance and error bars indicate ± standard deviation. Bars are colored according to ARG class and organized into 2 groups based on qualitative enrichment patterns: **A)** Beach to Ridge Gradient and **B)** Intermediate Enriched.

The isolate dataset had patterns that were broadly consistent with metagenomes (Figure 6). Isolate Clusters 1 and 2 exhibited significant negative correlations with the Physical Properties subgroup (Isolate Cluster 2: lm, p=0.018, R^2^=0.22, Figure 6A; Isolate Cluster 1: lm, p=0.014, R^2^=0.24, Figure 6E), significant positive correlations with the Enzymatic Activity group (Isolate Cluster 2: lm, p=0.01, R^2^=0.262, Figure 6B; Isolate Cluster 1: lm, p=0.002, R^2^=0.369, Figure 6F), as well as significant positive correlations with the Trace Elements group (Isolate Cluster 2: lm, p=0.004, R^2^=0.334, Figure 6C; Isolate Cluster 1: lm, p=0.003, R^2^=0.337, Figure 6G). These clusters included Vancomycin (Glycopeptide) resistance and Erythromycin (MLS) resistance that were significantly affected by site (Kruskal-Wallis Test, p<0.05, Figure 7), as well as other abundant isolate resistances including Ampicillin- and Rifamycin (Rifampin)-resistance. The combination of these relationships resulted in a significant effect of site on isolate cluster 1 (ANOVA, p=0.033, F=3.226, Figure 6H) and a similar pattern of beach-to-ridge gradient and intermediate site enrichment (Figure 6D and 6H) as was seen in the metagenomes.

**Figure 6:**
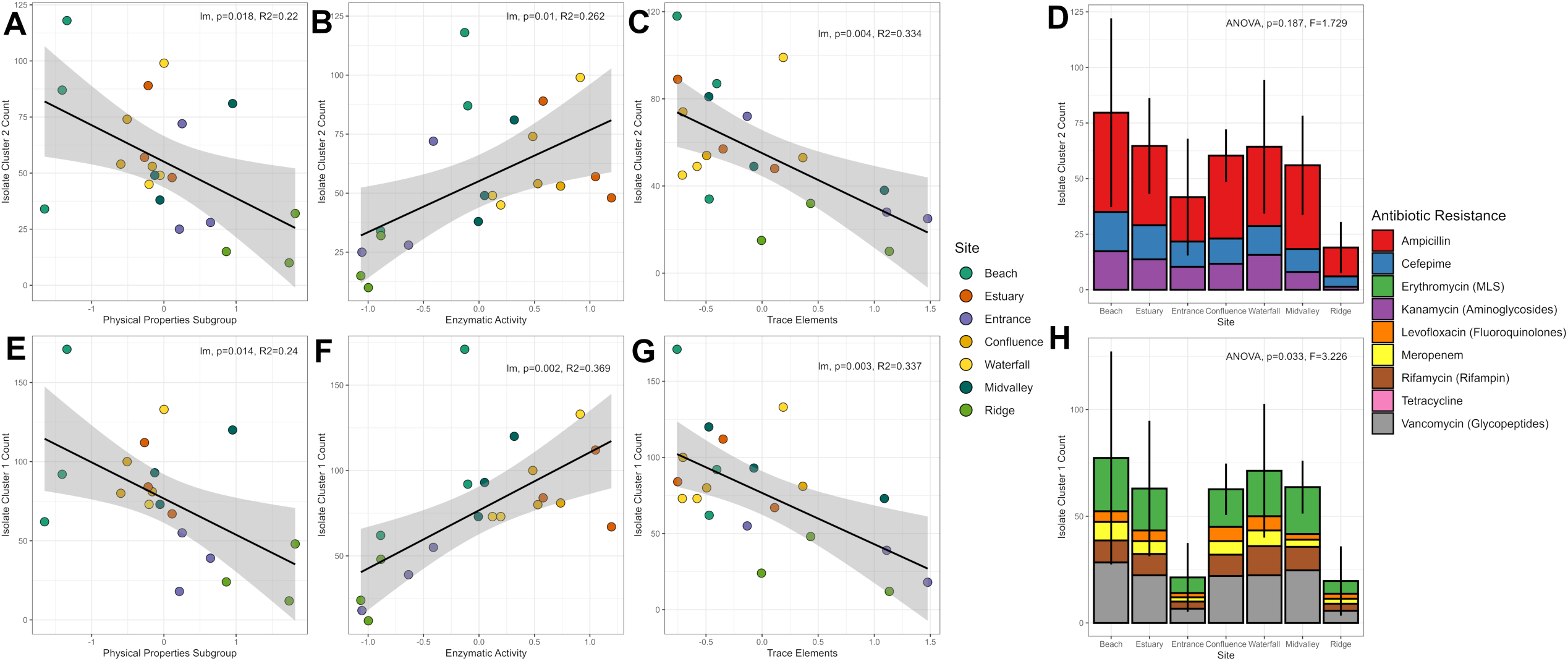
Significant correlations between soil properties and Isolate Clusters yield emergent patterns in the *in vitro* soil resistome across sites. Significant correlations (lm, p<0.05) between Isolate Cluster abundances and soil property groups defined in Figure 3 are displayed in the left 3 columns for **A-C)** Cluster 2 and **E-G)** Cluster 1. Indices for the soil groups were calculated by summing the z-score normalized values for every variable in a group. Resistant isolate counts of these clusters across sites are displayed in the righthand column for **D)** Cluster 2 and **H)** Cluster 1. Error bars indicate ± standard deviation, and bar colors indicate the resistance types in each cluster.

**Figure 7:**
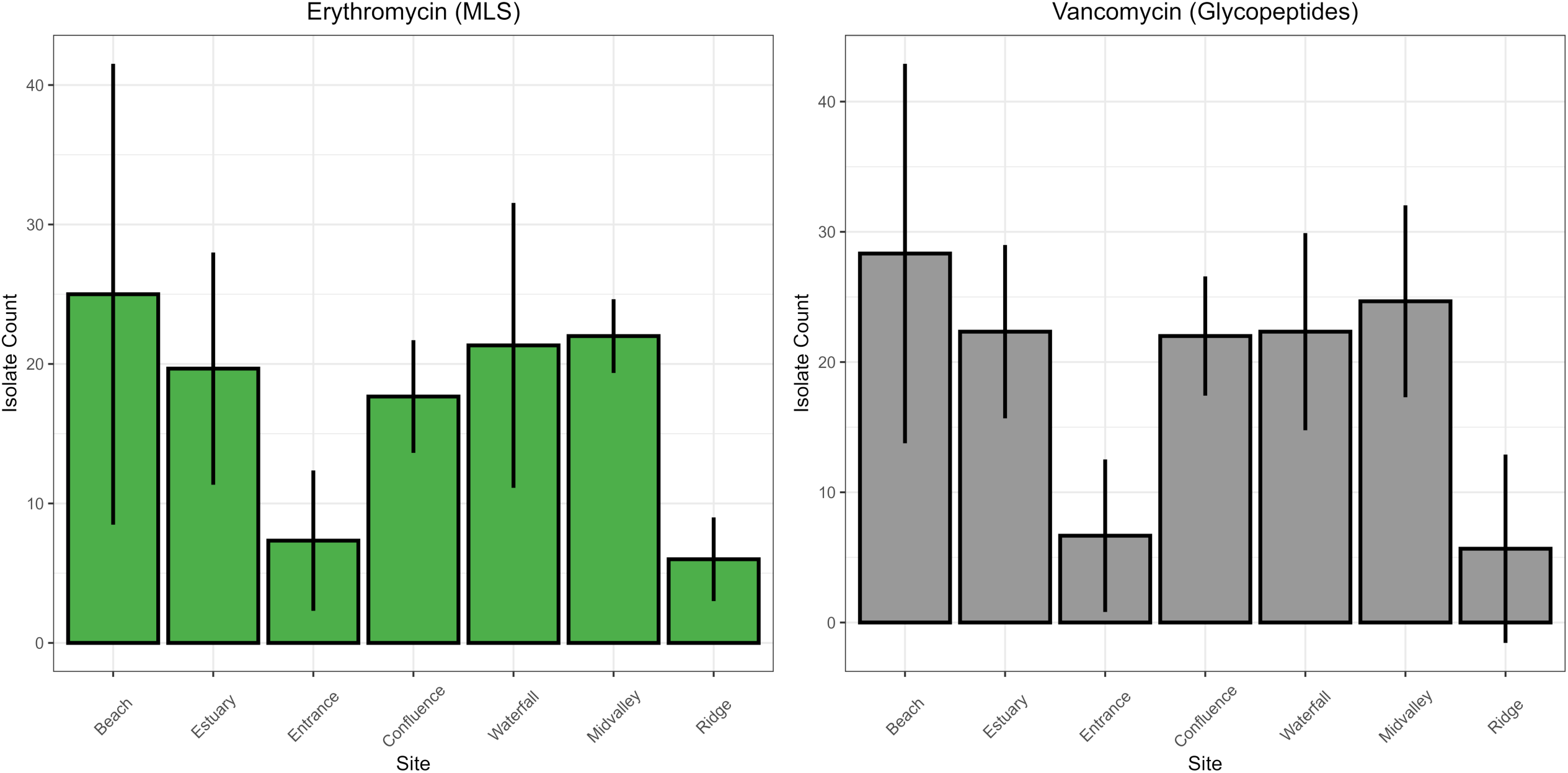
Isolate resistances that responded significantly to site (Kruskal-Wallis Test, p<0.05). Bars indicate mean isolate resistance counts and error bars indicate ± standard deviation. Plots are labeled according to resistance type. There were no significant pairwise differences between sites after p-value adjustment for multiple comparisons (Dunn’s test, adjusted-p > 0.05).

### Environmental Filtering of Resistome Hosts

Taxonomic composition of resistances, rather than functional identity, determined ARG and Isolate Cluster composition, linking specific bacterial taxa and their harbored resistances to gradients in soil properties. The taxonomic compositions of resistances, indicated by the relative abundances of bacterial classes assigned to each read/isolate (Figures 2 and 3), show clear distinctions between the various clusters. In the metagenomes, Actinomycetes were enriched in ARG Cluster 1, *Gammaproteobacteria* were more abundant in ARG Cluster 2, and *Betaproteobacteria* were enriched in ARG Cluster 3. In the isolates, Isolate Cluster 1 was enriched in *Gammaproteobacteria* whereas Isolate Cluster 2 had higher proportions of *Actinomycetes*. These qualitative patterns are corroborated when directly testing differences in taxonomic compositions between the various clusters (Figure 8). ARG taxonomic composition significantly correlated with cluster (PERMANOVA, R^2^=0.28516, F=10.107, p=0.001, Figure 8A), indicating that ARGs relation to soil properties was connected to their taxonomic composition. While resistant isolate taxonomic composition was not significantly correlated with cluster (PERMANOVA, R^2^=0.28402, F=1.1901, p=0.264, Figure 8B), cluster still explained a similar amount of variation compared to the metagenomes and was likely not significant due to the low sample size (n=9 resistance types) in the isolate dataset. In both datasets *Actinomycetes* and *Gammaproteobacteria* were significantly differentially abundant between the three clusters (ANOVA, p<0.05, Figure 8C). Furthermore, clusters that were enriched in these taxa exhibited consistent relationships with soil properties in both metagenomes and isolates (Figure 8D).

**Figure 8:**
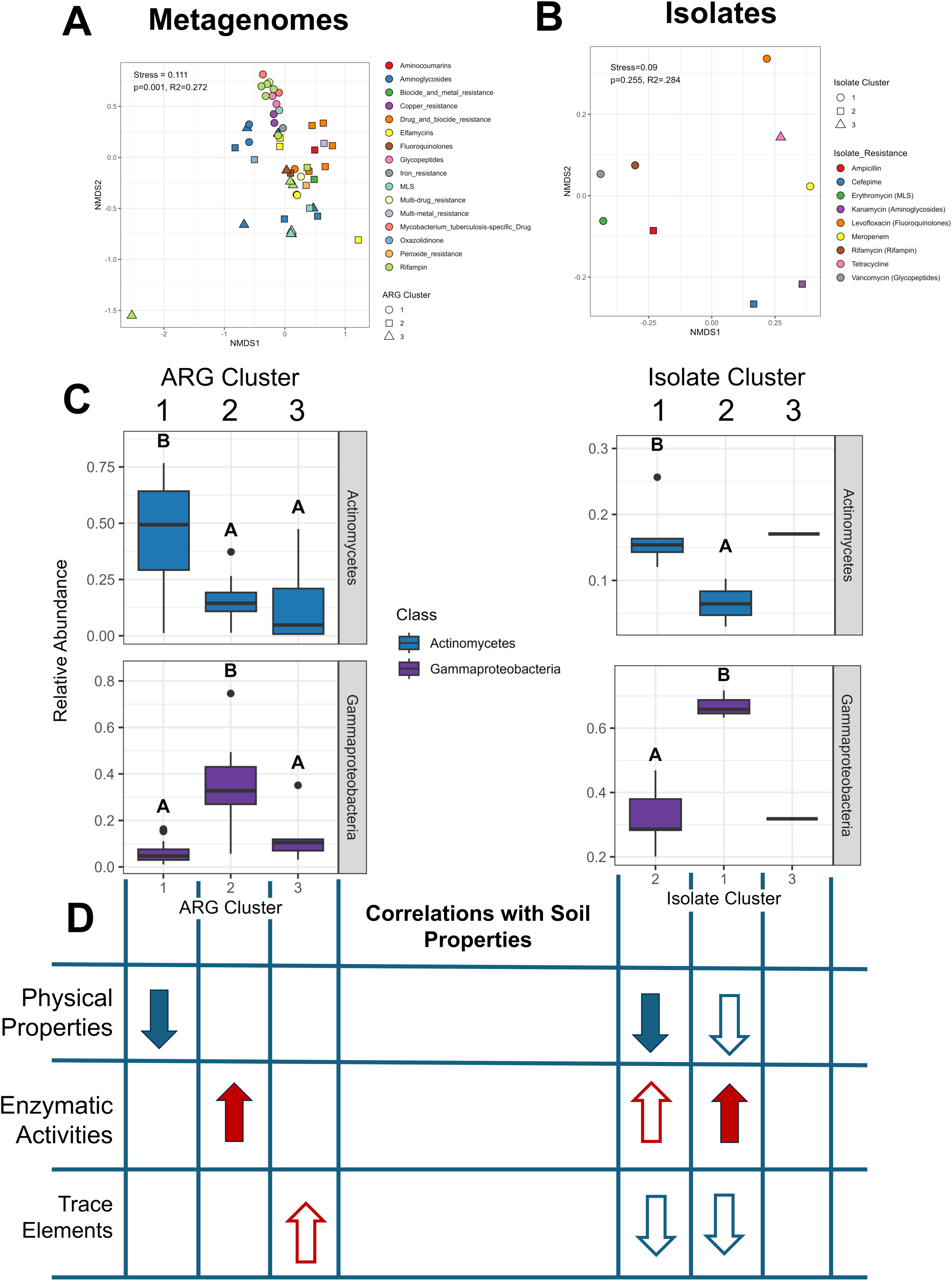
Taxonomic composition of soil resistomes. NMDS ordinations of the taxonomic composition of each resistance in **A)** metagenomes and **B)** isolates. Point shape corresponds to the cluster ID from Figures 2 and 3. P values and R^2^ values are presented from PERMANOVAs testing the effect of cluster ID on resistance taxonomic composition. **C)** Boxplots display bacterial classes shared in both resistome datasets that had significantly different enrichment between the 3 clusters (ANOVA, p<0.05). Letters denote significant pairwise differences (Tukey post hoc test, p<0.05). Isolate Cluster 3 had a single resistance and was excluded from pairwise significance testing. **D)** Schematic summarizing how the ARG/Isolate clusters (columns) correlated with the 3 main soil property groups (rows) from Figures 4 and 6. Arrows indicate a significant correlation, with arrow color and direction signifying the direction of correlation (red: positive, blue: negative). Filled arrows indicate that ARG clusters and Isolate clusters significantly enriched in the same bacterial class showed matching correlations with the same soil properties.

Specifically, clusters enriched in *Actinomycetes* exhibited significant negative correlations with Physical Properties and clusters enriched in *Gammaproteobacteria* exhibited significant correlations with Enzymatic Activities.

## DISCUSSION

In this study, we investigated drivers of the soil resistome across a tropical watershed with strong gradients in soil properties, integrating metagenomic analyses with culture-based validation. Broadly, we found that both metagenomic resistomes and isolate resistomes are influenced by sample site, but this is indirectly mediated via the interaction between resident bacterial communities and the unique soil properties at each site. Concordant patterns in these two resistome datasets are reflective of the unifying control of bacterial community structure on both *in situ* and *in vitro* resistomes.

### Bacterial Community Composition and Soil Properties Structures the Soil Resistome

Our combined results showed that bacterial community structure is an important determinant of the soil resistome. This suggests that resistome variation across the transect may result from the enrichment of distinct bacterial communities at the various sites. These findings reinforce other studies, which also found significant correlations between resistomes and resident bacterial community structure in a variety of environments including sewage, urban environments, and soils[10, 13, 42–45]. This concept counters the longstanding hypothesis that ARGs are uncoupled from microbial phylogenetic controls due to their high mobility^3,7,46,47^. It is worth noting that horizontal gene transfer, while possible between more distant relatives, more often occurs between similar species due to required alignments in recipient plasmids and cell wall proteins[48]. If horizontal gene transfer is phylogenetically restricted in this system, it could still play a major role in resistome structuring. Regardless of the exact mechanism, these data suggest that microbial phylogenetic identity may exert a larger degree of control on soil resistome structure than previously thought.

Specific soil properties, especially physical properties related to water dynamics, trace elements, and enzymatic activities, exhibited consistent patterns of correlation with resistance clusters in both datasets, indicating their importance in structuring the soil resistome *in situ* and *in vitro*. These properties have been previously shown to directly alter the soil resistome. High levels of soil moisture can inhibit the proliferation of ARGs from the environment[49] via reduction in cell membrane permeability and increased DNase activity[50], yielding negative correlations between soil water content and AMR in rice paddies[51] and other agricultural soils[52]. Various metals/trace elements have been shown to increase AMR abundances[12] via enhanced horizontal gene transfer from modulated conjugation[12, 53, 54], coselection with spatially colocalized ARGs[51], and induction of oxidative stress[49]. In addition, these soil properties have also been shown to exert strong influences on soil bacterial community structure more broadly[55–58], supporting our hypothesis that soil properties indirectly structure the soil resistome via influence on AMR-host distribution.

### Selection of Host Taxonomy by Soil Properties Indirectly Structures the Soil Resistome

The relationship between AMR and soil properties was driven primarily by resistance taxonomic composition rather than resistance function. This concept is supported by studies using Bayesian structural equation models, which have shown indirect effects of soil properties on the resistome via microbial host selection[10, 11, 59, 60]. However, few studies have validated modeling results with empirical correlations between soil properties and resistomes both *in situ* and *in vitro*.

Our findings add empirical evidence to the aforementioned conceptual model: ARG and isolate clusters that had distinct associations with soil properties exhibited unique taxonomic signatures, with *Actinomycetes*, *Betaproteobacteria*, and *Gammaproteobacteria* overrepresented in various resistance clusters. *Actinomycetes* was enriched in ARG Cluster 1 and are documented as abundant soil taxa adapted to low-moisture environments[61–63] that are inhibited under higher moisture conditions[64]. They also harbor diverse ARGs in both clinical and soil contexts, including vancomycin- and aminoglycoside-resistance genes consistent with those detected in this study[44, 49, 65–68]. The negative relationship between *Actinomycetes*-enriched ARGs and soil moisture in this watershed is therefore expected and follows a gradient of from the ridge, which has high water holding capacity and high gravimetric water content, to the beach with its substantially lower water holding capacity and gravimetric water content. *Betaproteobacteria* were associated with trace element-enriched soils and dominated Cluster 3 ARGs, including MLS- and rifampin-resistance genes enriched at intermediate sites. This pattern is consistent with previous reports that *Betaproteobacteria* harbor diverse resistances and increase in abundance with trace elements such as Cu, Zn, Mo, Mn, and Fe[55, 60]. Likewise, *Gammaproteobacteria* dominated ARG Cluster 2 and most cultured resistances and were positively associated with enzymatic activity metrics, aligning with their high metabolic activity and well-documented role as ARG hosts in soils and culture[45, 60, 69, 70]. Collectively, these data suggest a process by which soil properties indirectly shape the soil resistome via selection for bacterial taxa harboring unique AMR repertoires. Soil resistome structuring is therefore an indirect outcome of the larger environmental filtering of soil microbial communities.

### Parallel Patterns in Metagenomic- and Isolate-Datasets

Despite well-known differences between *in situ* molecular methods and culture-based approaches, our two datasets exhibited striking similarities. Differences likely arise from the widely documented distinctions between molecular- and culture-based methodologies. Amplicon data reflects the entire microbial community whereas culture data only reflects a subset that can be cultured[16]; metagenomes reflect functional potential whereas culture bioassays reflects actualized function^15,71^; and culture data only reflects microorganisms capable of growth under selected conditions. Additionally, certain phenotypes such as ampicillin (beta-lactamase)- and vancomycin (glycopeptides)-resistance involve multiple genes^74,75^, thus it is not easy to compare actualized drug resistance to *in situ* AMR genomic potential. Despite these differences, in this study we found consistent patterns between the metagenomic- and isoalte-resistomes. AMR abundance was significantly correlated between the two datasets, both resistomes were significantly correlated with bacterial community structure, and resistances enriched in the same taxonomic classes in the 2 datasets exhibited consistent correlations with the same soil properties. These consistencies indicate that *in situ* surveys of AMR genomic potential can be predictive of actualized AMR levels in culturable isolates from the same location.

The taxonomic patterns connected to ARGs were less pronounced in the isolate dataset, due in part to dominance of *Gammaproteobacteria* across most cultured resistances and potential bottlenecks of culturing fastidious bacterial taxa that are abundant in the metagenomes. Even though we used antibiotics in the initial isolation media for the cultures, we found that a number of strains (94, 10.9%) did not possess resistances in the downstream antibiotic susceptibility assays. While high proportions of antibiotic resistance in these isolates indicated that the selection did work, the number of isolates that lost their resistances through isolation may also indicate the transient nature of resistance (e.g. lost of resistant genes/plamids)[74]. This initial isolation approach could explain the remarkable consistency between the predicted metagenomic potential and the realized property of resistance in the isolates.

### Clinical and Landscape Implications

Identification of key soil properties and microbial taxa that influence resistomes at the landscape scale will allow researchers to a) generalize potential hotspot regions of high AMR and b) identify potential problematic taxa that carry clinically relevant AMR in soils. Based on this survey, physical properties altering water holding capacity and soil moisture, trace element concentrations, and enzymatic activities are potential key parameters to consider when examining environmental properties that structure the soil resistome. Although landscape-wide patterns are likely a result of soil property-AMR correlations, the abundance and diversity of AMR at any given site is the emergent result of multiple contrasting relationships between different soil properties and resistances and is thus difficult to predict *a priori*.

We also identified bacterial classes *Actinomycetes*, *Gammaproteobacteria*, and *Betaproteobacteria* that carried the majority of resistances and responded differentially to the above soil properties. The most abundant resistances in both datasets included clinically relevant resistances to Rifampin/Rifamycin, Glycopeptides/Vancomycin, Ampicillin, and MLS resistances, all of which are connected to taxa found in the WHO Bacterial Priority Pathogens 2024 List[75]. Among these, resistance to Vancomycin, a last-resort antibiotic, could lead to untreatable infections. The documented potential for these classes to carry, maintain, and even exchange ARGs with clinical pathogens[3] means these taxa may act as environmental AMR reservoirs for clinical pathogens.

Our direct antimicrobial resistance assays also revealed a high degree of multi-drug resistance (MDR) throughout the Waimea watershed, similar to observations in other soil environments[76–79]. In some cases, isolates had resistance to all 9 classes of antibiotics tested (Figure S7). These resistances may be attributed to efflux pumps found within the metagenomes throughout the transect. MDR has been found to be ubiquitous within the diverse bacterial communities of soils[78, 79] and appears ubiquitous across sites in this study, suggesting that the various sites and their associated soil properties had minimal effect on MDR distribution.

Of interest is the enrichment of many abundant, clinically relevant resistances at the lower/makai (ocean) sections of the watershed, especially the beach. Waimea Beach is a highly trafficked location for tourists and locals alike. While we do not have direct measurements of human visitations, daily visitors to the North Shore of O‘ahu top 12,000[80], with most of these visitors likely stopping at the iconic Waimea Beach. The decreasing gradient of AMR abundance in our transect tracks a decreasing human activity gradient from the beach to the extremely isolated ridge, which receives almost no human visitors due to its inaccessibility. The enrichment at more highly trafficked locations could be a result of multiple anthropogenic mechanisms such as horizontal gene transfer of resistances from the human microbiome to the environment, increased human-derived antibiotic load in the environment selecting for environmental AMR, and/or increased environmental toxins selecting for AMR[81]. However, this study was not designed to directly test these mechanisms, and we were unable to quantify human visitations at our sampling sites to calculate anthropogenic impacts. Parallel changes in both human activities and soil properties along the transect further obfuscate causal relationships between human activity and environmental AMR. To adequately assess the impact of human activity on soil AMR gradients, as well as enrichment leading to potential human infections[82], future studies should select sites with direct measures of human visitations while controlling for other key variables that potentially alter AMR in soils.

## CONCLUSIONS

Our study provides new insight into the ecological mechanisms shaping the soil resistome, a critical environmental reservoir of AMR with direct implications for public health. While AMR is a growing global threat, much remains unknown about how soil properties govern the distribution and abundance of ARGs in soil. By using both metagenomic and culture-based approaches across a geographically constrained transect with a gradient of soil properties, we show that the soil resistome structure is predominantly shaped by bacterial community composition, which is itself structured by soil properties. Key AMR-harboring taxa include *Actinomycetes*, *Gammaproteobacteria*, and *Betaproteobacteria*, which are correlated with soil physical properties, enzymatic activities, and trace elements, respectively. This suggests that resistome composition is a downstream consequence of environmentally structured microbial communities. These relationships result in concordant site-specific patterns in both metagenomic and culture datasets, particularly a decreasing abundance in common AMR types from the beach to the ridge. High AMR levels associated with human densities (such as the beach site in this study) deserves more focused studies. By linking soil physicochemistry, bacterial taxonomy, and resistance phenotypes within a single, ecologically coherent system, this study provides new insights for understanding how soil AMR emerges and persists in the environment, with direct relevance for AMR surveillance, management, and mitigation strategies.

## Availability of Data and Material

The datasets supporting the conclusions of this article are available in Sequence Read Archive (SRA) under project numbers PRJNA995197 (*in situ* 16S rRNA gene amplicons), PRJNA1463365 (cultured isolate 16S rRNA gene amplicons), and PRJNA1465928 (*in situ* metagenomes). All data used to generate figures has been deposited in GitHub at https://github.com/nnguyenlab/waimea-watershed-resistome and publicly accessioned via Zenodo (10.5281/zenodo.20190517).

## Acknowledgements

We thank our community partner, the Waimea Botanical Garden, especially Chad Middleton, for access to the garden and facilitating this work. We thank Erzsi Palko for assistance with part of the culturing work; Dr. Giovanna Slanzon, Jennifer Saito, and Dr. Karolina Peplowska for facilitating metagenomic sequencing; Dr. David Pompeani for the metals analysis; Dr. Enrique Doster and Nathalie Bonin for assistance with the AMR++ pipeline; and Kate Tracy for assistance with the graphical abstract. This project is part of a larger effort of the C-MĀIKI Institute to understand the connectivity of the microbiome to the land and water of an entire watershed, supported by the Office of the Vice Chancellor for Research at the University of Hawaici at Mānoa and the U.S. National Institutes of Health Center of Biomedical Research Excellence 1P20GM125508-01, project 5P20GM125508-03.

## Conflict of Interests

The authors declare that they have no competing interests.

## Ethics Statement

The content and authorship of the submitted manuscript have been approved by all authors, and that all prevailing local, national and international regulations and conventions, and normal scientific ethical practices, have been respected.

## Authors’ Contributions

Designed the project: NHN, SOIW, KKN Funding acquisition: NHN

Collected samples: SOIW, KKN

Processed samples: AL, CRF, CRH, JB, JND, MGT, RP, SL, FER Collected data: AL, ID, MTI

Analyzed data: WJS, SL, CRF, RP, TMM Performed visualizations: WJS, CRF, TMM

Writing – original draft: WJS, SL, NHN

Writing – review & editing: WJS, SL, MTI, AL, CRF, CRH, ID, JB, JND, RP, FER, MGT, SOIW, KKN, TMM, NHN

## Notes

### Competing Interest Statement

The authors have declared no competing interest.

## REFERENCES

1. O’Neill J. Tackling drug-resistant infections globally: final report and recommendations. Report. Government of the United Kingdom; 2016.

2. Organisation mondiale de la santé, editor. Antimicrobial resistance: global report on surveillance. Genève: World health organization; 2014.

3. Forsberg KJ, Reyes A, Wang B, Selleck EM, Sommer MOA, Dantas G. The Shared Antibiotic Resistome of Soil Bacteria and Human Pathogens. Science. 2012;337:1107–11. 10.1126/science.1220761.

4. Bottery MJ, Pitchford JW, Friman V-P. Ecology and evolution of antimicrobial resistance in bacterial communities. ISME J. 2021;15:939–48. 10.1038/s41396-020-00832-7.

5. Han B, Ma L, Yu Q, Yang J, Su W, Hilal MG, et al. The source, fate and prospect of antibiotic resistance genes in soil: A review. Front Microbiol. 2022;13:976657. 10.3389/fmicb.2022.976657.

6. Dantas G, Sommer MOA, Oluwasegun RD, Church GM. Bacteria Subsisting on Antibiotics. Science. 2008;320:100–3. 10.1126/science.1155157.

7. Wright GD. Antibiotic resistance in the environment: a link to the clinic? Curr Opin Microbiol. 2010;13:589–94. 10.1016/j.mib.2010.08.005.

8. Fierer N. Embracing the unknown: disentangling the complexities of the soil microbiome. Nat Rev Microbiol. 2017;15:579–90. 10.1038/nrmicro.2017.87.

9. Delgado-Baquerizo M, Hu H-W, Maestre FT, Guerra CA, Eisenhauer N, Eldridge DJ, et al. The global distribution and environmental drivers of the soil antibiotic resistome. Microbiome. 2022;10:219. 10.1186/s40168-022-01405-w.

10. Hu H, Wang J, Singh BK, Liu Y, Chen Y, Zhang Y, et al. Diversity of herbaceous plants and bacterial communities regulates soil resistome across forest biomes. Environ Microbiol. 2018;20:3186–200. 10.1111/1462-2920.14248.

11. Zhang Y, Cheng D, Zhang Y, Xie J, Xiong H, Wan Y, et al. Soil type shapes the antibiotic resistome profiles of long-term manured soil. Sci Total Environ. 2021;786:147361. 10.1016/j.scitotenv.2021.147361.

12. Shi X, Xia Y, Wei W, Ni B-J. Accelerated spread of antibiotic resistance genes (ARGs) induced by non-antibiotic conditions: Roles and mechanisms. Water Res. 2022;224:119060. 10.1016/j.watres.2022.119060.

13. Forsberg KJ, Patel S, Gibson MK, Lauber CL, Knight R, Fierer N, et al. Bacterial phylogeny structures soil resistomes across habitats. Nature. 2014;509:612–6. 10.1038/nature13377.

14. Li B, Yang Y, Ma L, Ju F, Guo F, Tiedje JM, et al. Metagenomic and network analysis reveal wide distribution and co-occurrence of environmental antibiotic resistance genes. ISME J. 2015;9:2490–502. 10.1038/ismej.2015.59.

15. Lewis WH, Tahon G, Geesink P, Sousa DZ, Ettema TJG. Innovations to culturing the uncultured microbial majority. Nat Rev Microbiol. 2020. 10.1038/s41579-020-00458-8.

16. Rappé MS, Giovannoni SJ. The Uncultured Microbial Majority. Annu Rev Microbiol. 2003;57:369–94. 10.1146/annurev.micro.57.030502.090759.

17. Amend AS, Swift SOI, Darcy JL, Belcaid M, Nelson CE, Buchanan J, et al. A ridge-to-reef ecosystem microbial census reveals environmental reservoirs for animal and plant microbiomes. Proc Natl Acad Sci. 2022;119:e2204146119. 10.1073/pnas.2204146119.

18. Hynson NA, Frank KL, Alegado RA, Amend AS, Arif M, Bennett GM, et al. Synergy among Microbiota and Their Hosts: Leveraging the Hawaiian Archipelago and Local Collaborative Networks To Address Pressing Questions in Microbiome Research. mSystems. 2018;3. 10.1128/mSystems.00159-17.

19. Hynson NA, Frank KL, Alegado RA, Amend AS, Arif M, Bennett GM, et al. Synergy among Microbiota and Their Hosts: Leveraging the Hawaiian Archipelago and Local Collaborative Networks To Address Pressing Questions in Microbiome Research. mSystems. 2018;3. 10.1128/mSystems.00159-17.

20. Crow SE, Hubanks H, Deenik JL, McClellan Maaz T, Tallamy Glazer C, Vizka E, et al. The legacy of intensive agricultural history on the soil health of (sub)tropical landscapes. Front Environ Sci. 2023;10. 10.3389/fenvs.2022.991262.

21. Maaz TM, Heck RH, Glazer CT, Loo MK, Zayas JR, Krenz A, et al. Measuring the immeasurable: A structural equation modeling approach to assessing soil health. Sci Total Environ. 2023;870:161900. 10.1016/j.scitotenv.2023.161900.

22. Kirby-Bauer-Disk-Diffusion-Susceptibility-Test-Protocol-pdf.

23. Heisey S, Ryals R, Maaz TM, Nguyen NH. A Single Application of Compost Can Leave Lasting Impacts on Soil Microbial Community Structure and Alter Cross-Domain Interaction Networks. Front Soil Sci. 2022;2. 10.3389/fsoil.2022.749212.

24. Apprill A, McNally S, Parsons R, Weber L. Minor revision to V4 region SSU rRNA 806R gene primer greatly increases detection of SAR11 bacterioplankton. Aquat Microb Ecol. 2015;75:129–37. 10.3354/ame01753.

25. Parada AE, Needham DM, Fuhrman JA. Every base matters: assessing small subunit rRNA primers for marine microbiomes with mock communities, time series and global field samples. Environ Microbiol. 2016;18:1403–14. 10.1111/1462-2920.13023.

26. Nguyen NH, Smith D, Peay K, Kennedy P. Parsing ecological signal from noise in next generation amplicon sequencing. New Phytol. 2015;205:1389–93. 10.1111/nph.12923.

27. Nguyen N, Lee S, Lin A. High-throughput cultivation and identification of soil bacteria. 2024.

28. Quince C, Lanzen A, Davenport RJ, Turnbaugh PJ. Removing Noise From Pyrosequenced Amplicons. BMC Bioinformatics. 2011;12:38. 10.1186/1471-2105-12-38.

29. Bolyen E, Rideout JR, Dillon MR, Bokulich NA, Abnet CC, Al-Ghalith GA, et al. Reproducible, interactive, scalable and extensible microbiome data science using QIIME 2. Nat Biotechnol. 2019;37:852–7. 10.1038/s41587-019-0209-9.

30. Schloss PD. Amplicon Sequence Variants Artificially Split Bacterial Genomes into Separate Clusters. mSphere. 2021;6:. 10.1128/msphere.00191-21.

31. Bonin N, Doster E, Worley H, Pinnell LJ, Bravo JE, Ferm P, et al. MEGARes and AMR++, v3.0: an updated comprehensive database of antimicrobial resistance determinants and an improved software pipeline for classification using high-throughput sequencing. Nucleic Acids Res. 2023;51:D744–52. 10.1093/nar/gkac1047.

32. Bolger AM, Lohse M, Usadel B. Trimmomatic: a flexible trimmer for Illumina sequence data. Bioinformatics. 2014;30:2114–20. 10.1093/bioinformatics/btu170.

33. Lakin SM, Dean C, Noyes NR, Dettenwanger A, Ross AS, Doster E, et al. MEGARes: an antimicrobial resistance database for high throughput sequencing. Nucleic Acids Res. 2017;45 Database issue:D574–80. 10.1093/nar/gkw1009.

34. Li H, Durbin R. Fast and accurate short read alignment with Burrows–Wheeler transform. Bioinformatics. 2009;25:1754–60. 10.1093/bioinformatics/btp324.

35. Wood DE, Lu J, Langmead B. Improved metagenomic analysis with Kraken 2. Genome Biol. 2019;20:257. 10.1186/s13059-019-1891-0.

36. R Core Team. R: A language and environment for statistical computing. 2013.

37. Oksanen J. Vegan: an introduction to ordination. 2013.

38. Breiman L, Cutler A, Liaw A, Wiener M. randomForest: Breiman and Cutlers Random Forests for Classification and Regression. 2024.

39. Fox J, Marquez M. RcmdrMisc: R Commander Miscellaneous Functions. 2014;:2.9–2. 10.32614/CRAN.package.RcmdrMisc.

40. Gu Z. Complex heatmap visualization. iMeta. 2022;1:e43. 10.1002/imt2.43.

41. Benjamini Y, Hochberg Y. Controlling the False Discovery Rate: A Practical and Powerful Approach to Multiple Testing. J R Stat Soc Ser B Methodol. 1995;57:289–300.

42. Gibson MK, Forsberg KJ, Dantas G. Improved annotation of antibiotic resistance determinants reveals microbial resistomes cluster by ecology. ISME J. 2015;9:207–16. 10.1038/ismej.2014.106.

43. Pehrsson EC, Tsukayama P, Patel S, Mejía-Bautista M, Sosa-Soto G, Navarrete KM, et al. Interconnected microbiomes and resistomes in low-income human habitats. Nature. 2016;533:212–6. 10.1038/nature17672.

44. Qian X, Gunturu S, Guo J, Chai B, Cole JR, Gu J, et al. Metagenomic analysis reveals the shared and distinct features of the soil resistome across tundra, temperate prairie, and tropical ecosystems. Microbiome. 2021;9:108. 10.1186/s40168-021-01047-4.

45. Su J-Q, Wei B, Ou-Yang W-Y, Huang F-Y, Zhao Y, Xu H-J, et al. Antibiotic Resistome and Its Association with Bacterial Communities during Sewage Sludge Composting. Environ Sci Technol. 2015;49:7356–63. 10.1021/acs.est.5b01012.

46. Smillie CS, Smith MB, Friedman J, Cordero OX, David LA, Alm EJ. Ecology drives a global network of gene exchange connecting the human microbiome. Nature. 2011;480:241–4. 10.1038/nature10571.

47. Stokes HW, Gillings MR. Gene flow, mobile genetic elements and the recruitment of antibiotic resistance genes into Gram-negative pathogens. FEMS Microbiol Rev. 2011;35:790–819. 10.1111/j.1574-6976.2011.00273.x.

48. Redondo-Salvo S, Fernández-López R, Ruiz R, Vielva L, De Toro M, Rocha EPC, et al. Pathways for horizontal gene transfer in bacteria revealed by a global map of their plasmids. Nat Commun. 2020;11:3602. 10.1038/s41467-020-17278-2.

49. Wang L, Gu X, Hui K, Yu T, Yuan Y, Chen G, et al. Interactions between antibiotic resistance genes and soil environmental factors: Coupling, antagonism, and synergism. Emerg Contam. 2025;11:100578. 10.1016/j.emcon.2025.100578.

50. Kittredge HA, Dougherty KM, Evans SE. Dead but Not Forgotten: How Extracellular DNA, Moisture, and Space Modulate the Horizontal Transfer of Extracellular Antibiotic Resistance Genes in Soil. Appl Environ Microbiol. 2022;88:e02280–21. 10.1128/aem.02280-21.

51. Han B, Yang F, Shen S, Mu M, Zhang K. Effects of soil habitat changes on antibiotic resistance genes and related microbiomes in paddy fields. Sci Total Environ. 2023;895:165109. 10.1016/j.scitotenv.2023.165109.

52. Zhou Y, Niu L, Zhu S, Lu H, Liu W. Occurrence, abundance, and distribution of sulfonamide and tetracycline resistance genes in agricultural soils across China. Sci Total Environ. 2017;599–600:1977–83. 10.1016/j.scitotenv.2017.05.152.

53. Lin H, Jiang L, Li B, Dong Y, He Y, Qiu Y. Screening and evaluation of heavy metals facilitating antibiotic resistance gene transfer in a sludge bacterial community. Sci Total Environ. 2019;695:133862. 10.1016/j.scitotenv.2019.133862.

54. Zhang Y, Gu AZ, Cen T, Li X, He M, Li D, et al. Sub-inhibitory concentrations of heavy metals facilitate the horizontal transfer of plasmid-mediated antibiotic resistance genes in water environment. Environ Pollut. 2018;237:74–82. 10.1016/j.envpol.2018.01.032.

55. Dai Z, Guo X, Lin J, Wang X, He D, Zeng R, et al. Metallic micronutrients are associated with the structure and function of the soil microbiome. Nat Commun. 2023;14:8456. 10.1038/s41467-023-44182-2.

56. Peng Z, Liang C, Gao M, Qiu Y, Pan Y, Gao H, et al. The neglected role of micronutrients in predicting soil microbial structure. Npj Biofilms Microbiomes. 2022;8:103. 10.1038/s41522-022-00363-3.

57. Schimel JP. Life in Dry Soils: Effects of Drought on Soil Microbial Communities and Processes. Annu Rev Ecol Evol Syst. 2018;49 Volume 49, 2018:409–32. 10.1146/annurev-ecolsys-110617-062614.

58. Shawver S, Ishii S, Strickland MS, Badgley B. Soil type and moisture content alter soil microbial responses to manure from cattle administered antibiotics. Environ Sci Pollut Res. 2024;31:27259–72. 10.1007/s11356-024-32903-z.

59. Yan Z-Z, Chen Q-L, Li C-Y, Thi Nguyen B-A, Zhu Y-G, He J-Z, et al. Biotic and abiotic factors distinctly drive contrasting biogeographic patterns between phyllosphere and soil resistomes in natural ecosystems. ISME Commun. 2021;1:13. 10.1038/s43705-021-00012-4.

60. Zheng D, Yin G, Liu M, Hou L, Yang Y, Van Boeckel TP, et al. Global biogeography and projection of soil antibiotic resistance genes. Sci Adv. 2022;8:eabq8015. 10.1126/sciadv.abq8015.

61. Williams ST, Shameemullah M, Watson ET, Mayfield CI. Studies on the ecology of actinomycetes in soil—VI. The influence of moisture tension on growth and survival. Soil Biol Biochem. 1972;4:215–25. 10.1016/0038-0717(72)90014-4.

62. Zvyagintsev DG, Zenova GM, Sudnizin II, Doroshenko EA. The Ability of Soil Actinomycetes to Develop at an Extremely Low Humidity. Dokl Biol Sci. 2005;405:461–3. 10.1007/s10630-005-0165-z.

63. Zvyagintsev DG, Zenova GM, Doroshenko EA, Gryadunova AA, Gracheva TA, Sudnitsyn II. Actinomycete growth in conditions of low moisture. Biol Bull. 2007;34:242–7. 10.1134/S1062359007030053.

64. Borowik A, Wyszkowska J. Soil moisture as a factor affecting the microbiological and biochemical activity of soil. Plant Soil Environ. 2016;62:250–5. 10.17221/158/2016-PSE.

65. Rahdar HA, Mahmoudi S, Bahador A, Ghiasvand F, Sadeghpour Heravi F, Feizabadi MM. Molecular identification and antibiotic resistance pattern of actinomycetes isolates among immunocompromised patients in Iran, emerging of new infections. Sci Rep. 2021;11:10745. 10.1038/s41598-021-90269-5.

66. Yushchuk O, Binda E, Marinelli F. Glycopeptide Antibiotic Resistance Genes: Distribution and Function in the Producer Actinomycetes. Front Microbiol. 2020;11. 10.3389/fmicb.2020.01173.

67. Yi X, Liang J-L, Su J-Q, Jia P, Lu J, Zheng J, et al. Globally distributed mining-impacted environments are underexplored hotspots of multidrug resistance genes. ISME J. 2022;16:2099–113. 10.1038/s41396-022-01258-z.

68. Liao H, Wen C, Huang D, Liu C, Gao T, Du Q, et al. Harnessing phage consortia to mitigate the soil antibiotic resistome by targeting keystone taxa Streptomyces. Microbiome. 2025;13:127. 10.1186/s40168-025-02117-7.

69. Esiobu N, Armenta L, Ike J. Antibiotic resistance in soil and water environments. Int J Environ Health Res. 2002;12:133–44. 10.1080/09603120220129292.

70. Kurm V, van der Putten WH, de Boer W, Naus-Wiezer S, Hol WHG. Low abundant soil bacteria can be metabolically versatile and fast growing. Ecology. 2017;98:555–64. 10.1002/ecy.1670.

71. Handelsman J. Metagenomics: Application of Genomics to Uncultured Microorganisms. Microbiol Mol Biol Rev. 2004;68:669–85. 10.1128/mmbr.68.4.669-685.2004.

72. Poole K. Resistance to β-lactam antibiotics. Cell Mol Life Sci CMLS. 2004;61:2200–23. 10.1007/s00018-004-4060-9.

73. Stogios PJ, Savchenko A. Molecular mechanisms of vancomycin resistance. Protein Sci. 2020;29:654–69. 10.1002/pro.3819.

74. Bag A, Kumar V, Adhikari A, Mandal B, Dhar S, Das BK. Impact of repeated in-vitro bacterial culture on virulence and antibiotic resistance characteristics: a study of Gram-positive and Gram-negative fish pathogens. Front Microbiol. 2025;16. 10.3389/fmicb.2025.1601681.

75. WHO Bacterial Priority Pathogens List 2024: Bacterial Pathogens of Public Health Importance, to Guide Research, Development, and Strategies to Prevent and Control Antimicrobial Resistance. 1st ed. Geneva: World Health Organization; 2024.

76. Bhullar K, Waglechner N, Pawlowski A, Koteva K, Banks ED, Johnston MD, et al. Antibiotic Resistance Is Prevalent in an Isolated Cave Microbiome. PLOS ONE. 2012;7:e34953. 10.1371/journal.pone.0034953.

77. Brown MG, Balkwill DL. Antibiotic Resistance in Bacteria Isolated from the Deep Terrestrial Subsurface. Microb Ecol. 2009;57:484–93. 10.1007/s00248-008-9431-6.

78. Van Goethem MW, Pierneef R, Bezuidt OKI, Van De Peer Y, Cowan DA, Makhalanyane TP. A reservoir of ‘historical’ antibiotic resistance genes in remote pristine Antarctic soils. Microbiome. 2018;6:40. 10.1186/s40168-018-0424-5.

79. Walsh F, Duffy B. The Culturable Soil Antibiotic Resistome: A Community of Multi-Drug Resistant Bacteria. PLOS ONE. 2013;8:e65567. 10.1371/journal.pone.0065567.

80. Kamita RY. Visitors to the North Shore. Research and Economic Analysis Division Department of Business, Economic Development & Tourism State of Hawaii; 2024.

81. Wang X, Chen Z, Mu Q, Wu X, Zhang J, Mao D, et al. Ionic Liquid Enriches the Antibiotic Resistome, Especially Efflux Pump Genes, Before Significantly Affecting Microbial Community Structure. Environ Sci Technol. 2020;54:4305–15. 10.1021/acs.est.9b04116.

82. Zhao Y, Li L, Huang Y, Xu X, Liu Z, Li S, et al. Global soil antibiotic resistance genes are associated with increasing risk and connectivity to human resistome. Nat Commun. 2025;16:7141. 10.1038/s41467-025-61606-3.

