## Supplementary Figures for "Soil Resistomes in a Tropical Watershed are Indirectly Structured by Bacterial Community Interactions with Soil Properties"

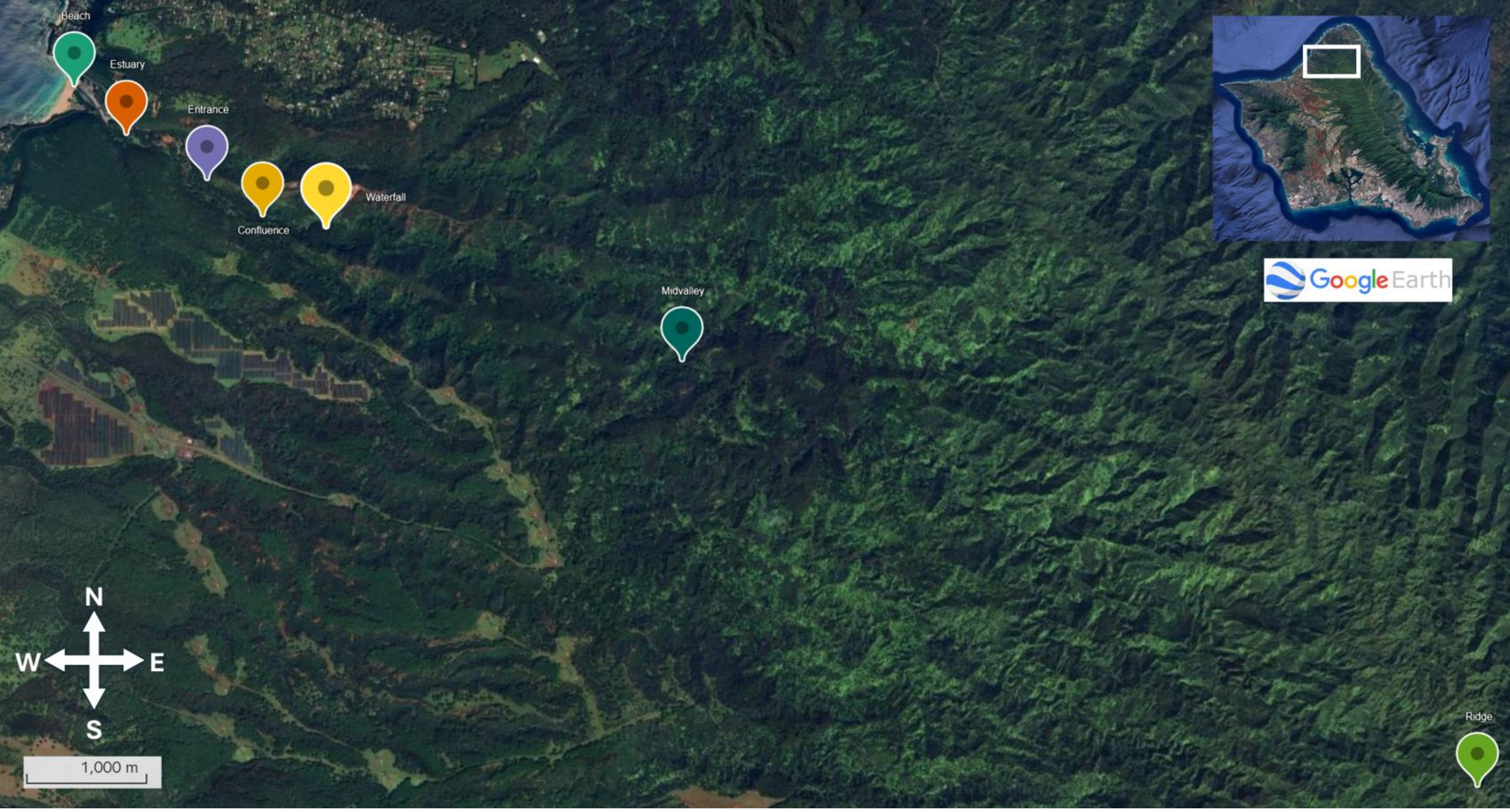


**Figure S1**: Sample sites of the transect of Waimea Valley on the north shore of O‘ahu, Hawai‘i

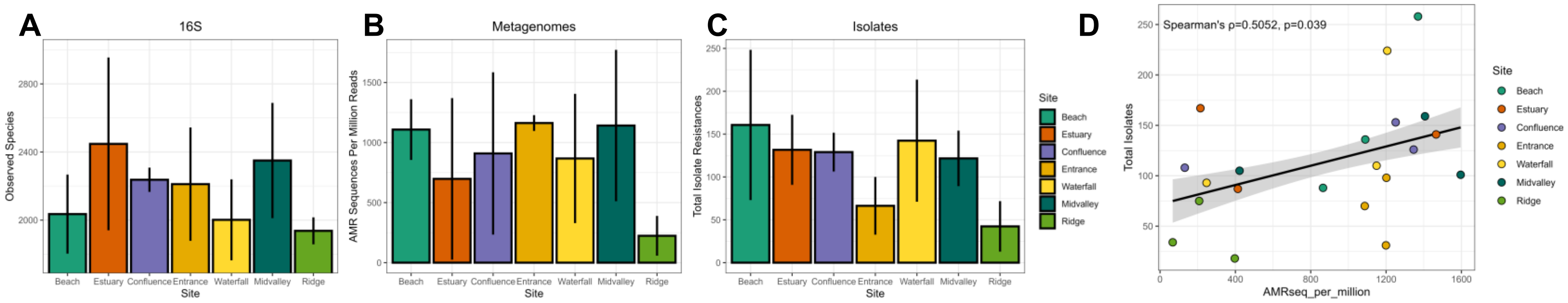


**Figure S2**: Richness across the surveyed sites for **A)** observed bacterial taxa in the 16S rDNA dataset, **B)** normalized AMR sequences in the metagenomes, and **C)** total number of antibiotic resistances in the isolate dataset. Bars are colored according to Site and error bars indicate ± standard deviation. **D)** Total isolate resistances significantly positively correlated with AMR sequence abundance in the metagenomes.

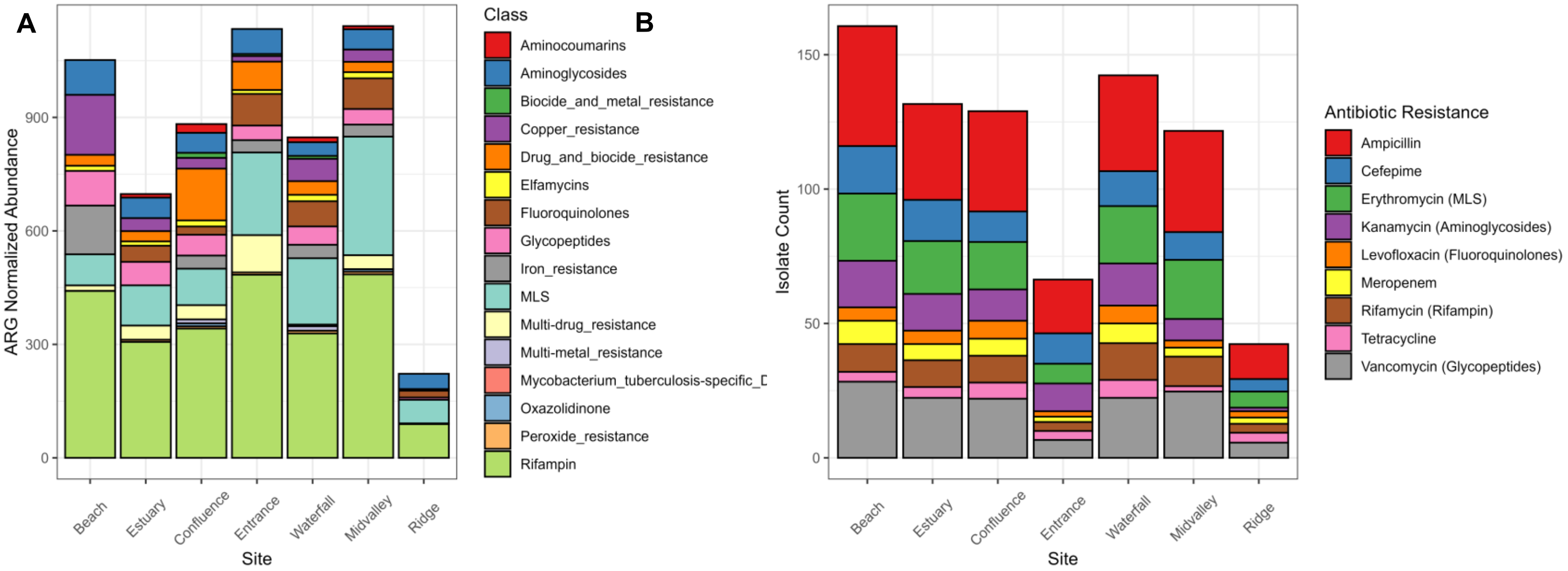


**Figure S3:** Distribution of AMR types across the surveyed sites for **A**) metagenomes (mean normalized ARG abundances) and **B**) isolates (mean resistant isolate counts). Colors indicate resistance type at the ARG class level for metagenomes and the resistance assay type for isolates.

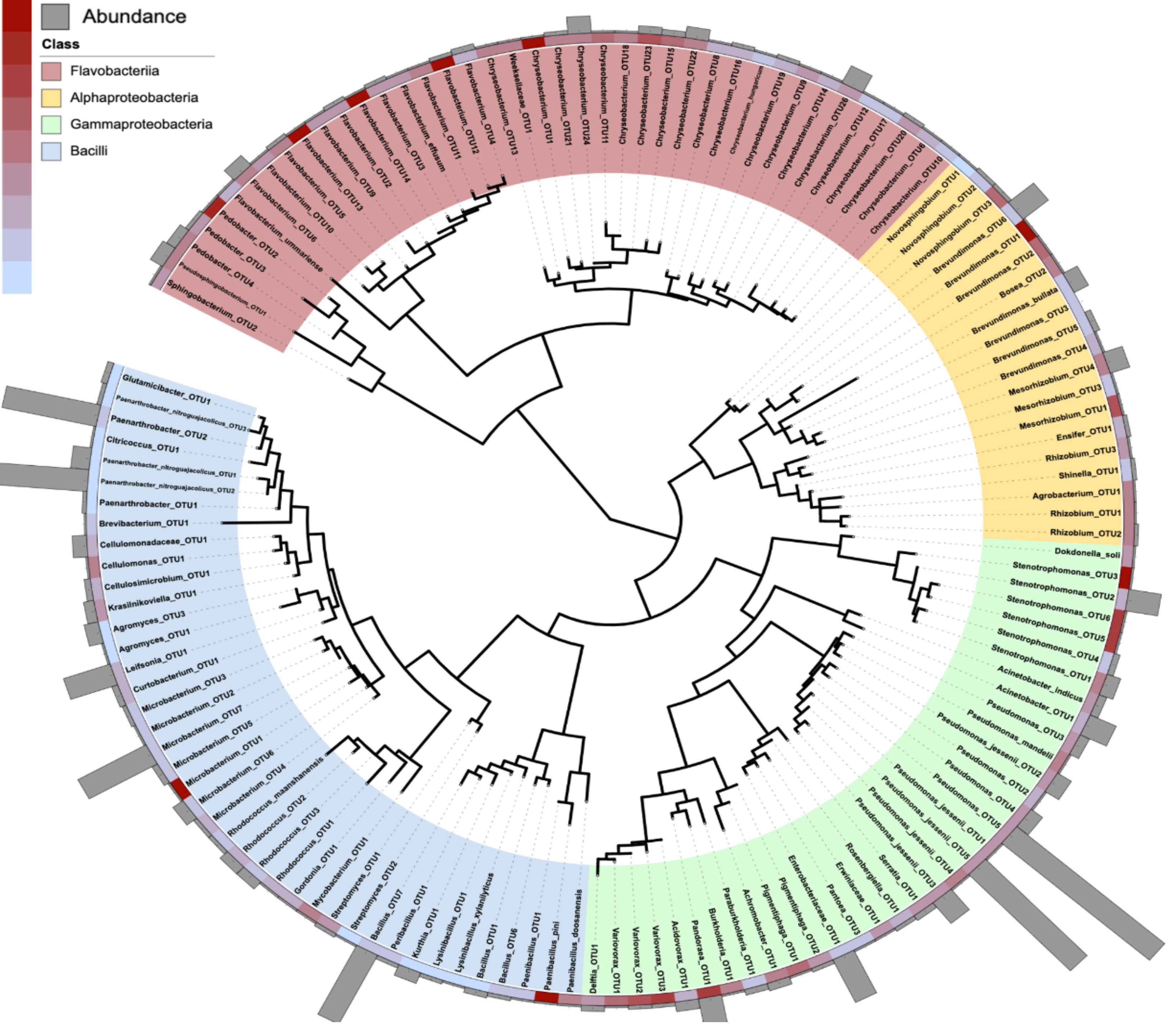


**Figure S4:** Phylogenetic tree of all cultured isolates, identified to the genus level. Inner ring 1 shows abundance of isolate; Inner ring 2 shows the levels of resistance from no resistance (blue) to high (red). Outer ring shows isolates that have 100% match to community data.

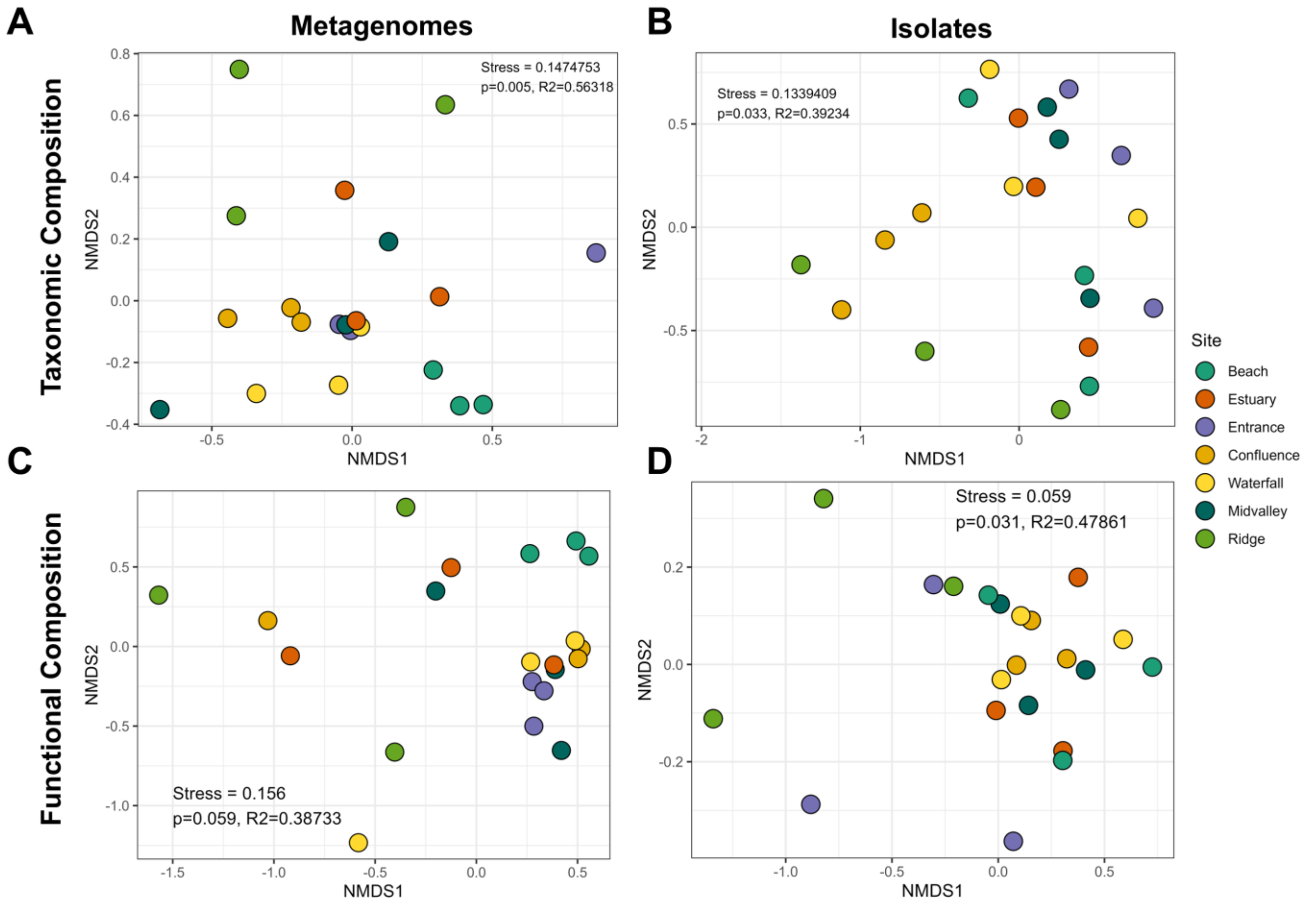


**Figure S5:** NMDS of resistome taxonomic and functional compositions by site in **A**) and **C**) metagenomes and **B**) and **D**) isolates. P values and $R^2$ values are presented from PERMANOVAs testing the effect of site on resistomes taxonomic and functional compositions.

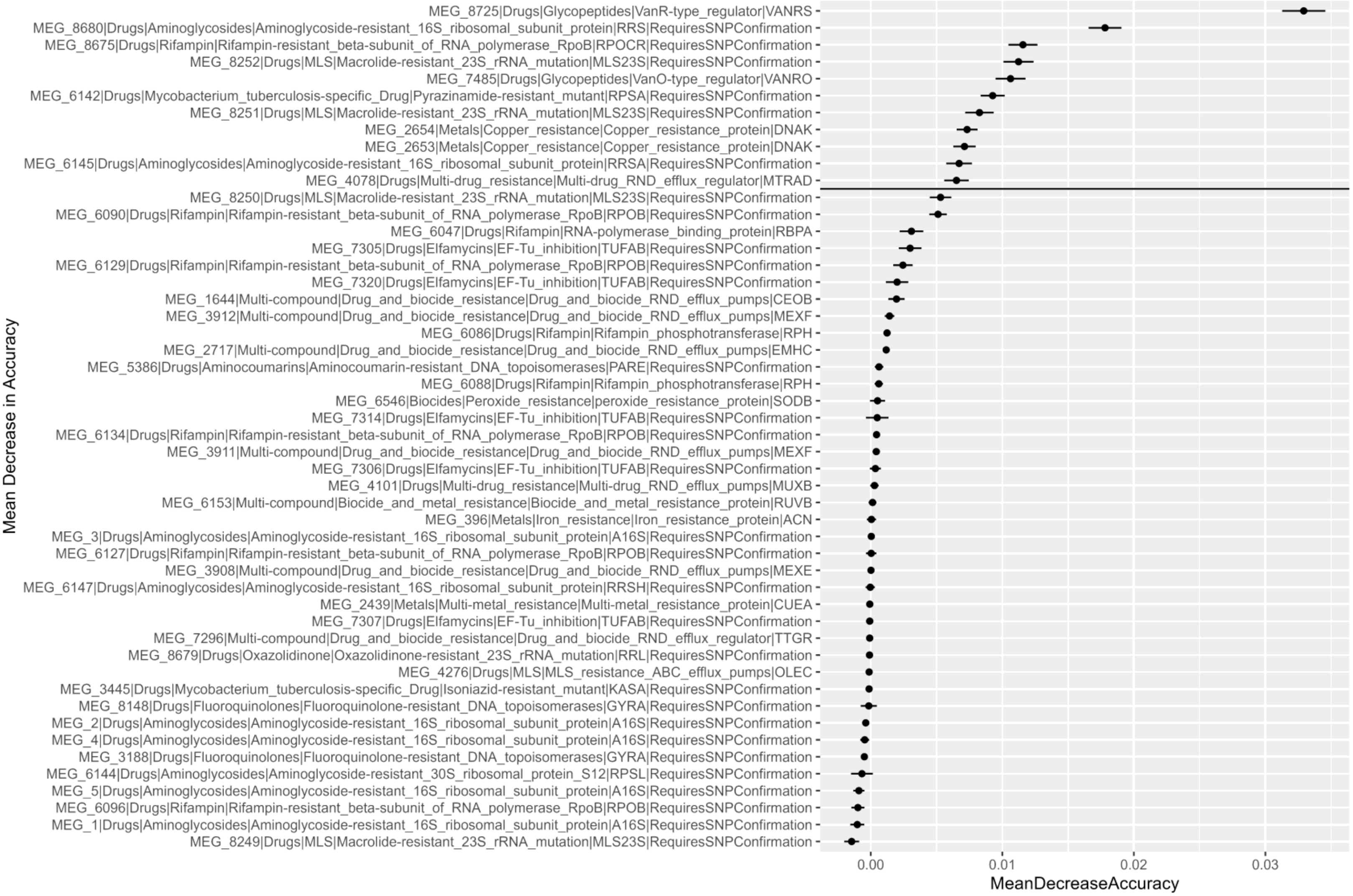


**Figure S6**: Variable importance plot of mean decrease in accuracy (MDA) values for all ARGs tested as predictors for survey site. Error bars indicate MDA value standard deviations. Horizontal line separates ARGs with MDA > 0.006.

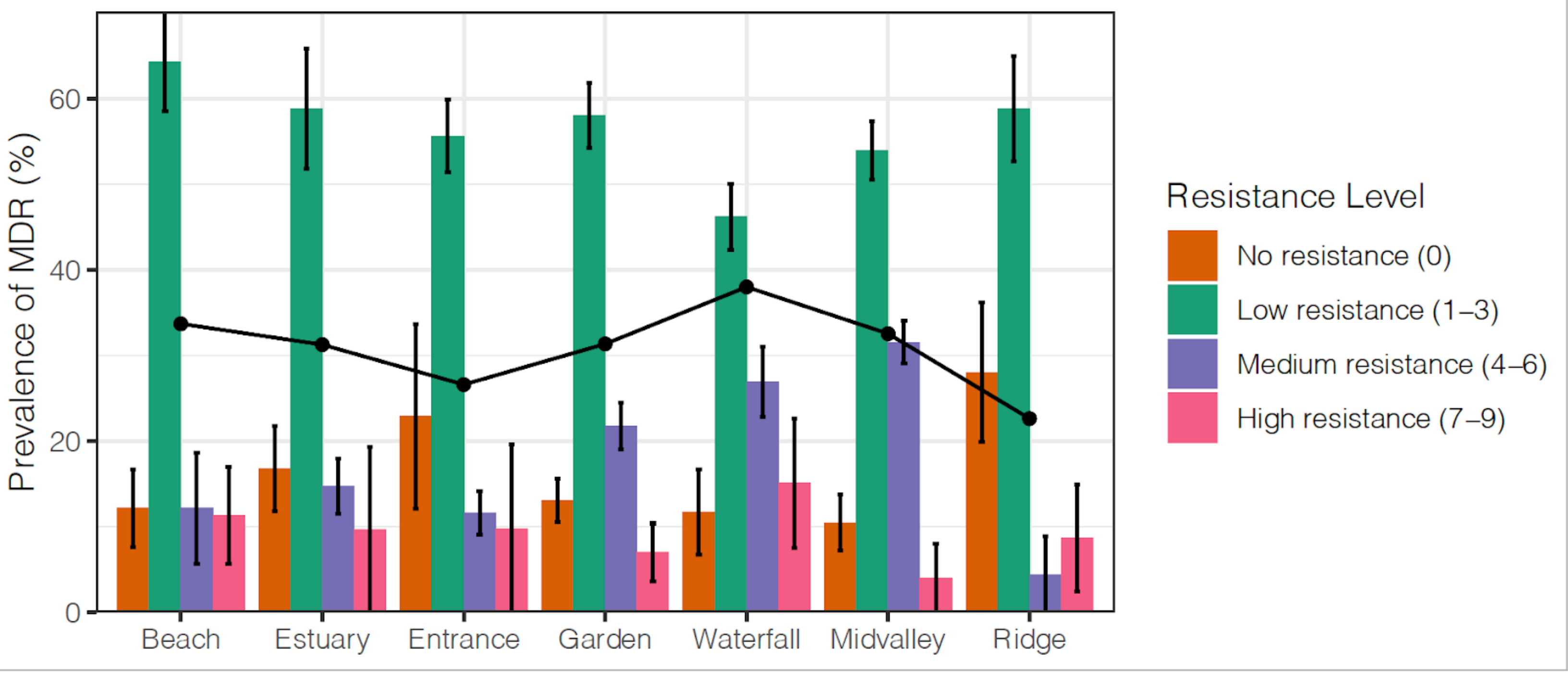


**Figure S7:** Prevalence of MDR across seven sites in the Waimea watershed. The prevalence of MDR was calculated at each site and normalized by dividing by the total number of cultured isolates at that site, regardless of their MDR. MDR was unambiguously grouped into no (0) low (1-3), medium (4-6), and high (7-9) MDR. Error bars represent the standard error of the mean.
