## Supplementary Methods for "Soil Resistomes in a Tropical Watershed are Indirectly Structured by Bacterial Community Interactions with Soil Properties"

**Bacteria isolation and antibiotic susceptibility test**

Isolation: The antibiotic concentration used was determined by minimum inhibitory concentration (MIC) for various microbial isolates. A slurry was prepared by vortexing 1 g of homogenized soil with 5 mL of sterile water, followed by serial dilution to 10⁻⁴, 10⁻⁵, and 10⁻⁶. All three dilutions were plated on each antibiotic selection medium. Plates were incubated at 25 °C for 3-7 days prior to picking morphologically different colonies from all dilutions.

Assay: Each isolate was grown in 5 mL of LB broth and incubated in a shaker at 37 °C for 16-24 hours. Once turbid, 300 µL was spread evenly onto a plate containing 30 ml (4 mm deep) Mueller-Hinton agar. Antibiotic disks (6.5 mm) containing standard concentrations of the antibiotic Ampicillin (10 µg), Cefepime (30 µg), Erythromycin (15 µg), Kanamycin (30 µg), Levofloxacin (5 µg), Meropenem (10 µg), Rifamycin (5 µg), Tetracycline (30 µg), Vancomycin (30 µg) were placed in the middle of the plate. A disk without antibiotics was used as a negative control. Plates were incubated overnight at 37 °C. The area of inhibited bacterial growth was measured and recorded (Table S3).

**DNA Extraction and Sequencing Methods**

Soil 16S PCR: First PCR parameters: polymerase activation at 98 °C for 30 s, followed by 20 cycles of 98 °C for 10 s, 55 °C for 15 s, 72 °C for 10 s, and a final extension at 72 °C for 7 minutes. The second PCR parameters were: polymerase activation at 98°C for 30 s, followed by 15 cycles of 98 °C for 10 s, 52 °C for 15 s, 72 °C for 10 s, and a final extension at 72 °C for 7 minutes.

Isolate Crude Extraction and 16S PCR: Crude extracts were collected by mixing 10 µl of cells with 100 µl alkaline lysis buffer (0.2 M Tris pH8, 0.011 M EDTA disodium, 0.25 M KCl, titrated with 1M NaOH to pH 10), incubated at 25 °C for 10 min followed by 95 °C for 10 min, and neutralized with 100 µl of neutralization solution (3% BSA w/v, 1 mM MgCl_2_) in 96-well plates. The plate was centrifuged for 5 min at 4000 ྾ g to pellet the cells. PCR parameters were the same for both PCR steps: polymerase activation at 95 °C for 60 s, followed by 34 cycles of 95 °C for 15 s, 53 °C for 15 s, 72 °C for 15 s, and a final extension at 72 °C for 7 min.

**Bioinformatics**

Soil 16S Bioinformatics: Demultiplexed sequences were imported into QIIME2 where primers, indexes, and adapters were removed using the CUTADAPT plugin^30^ leaving only the targeted sequence. Sequences were then trimmed based on quality, paired, and denoised using the DADA2 plugin^31^, followed by taxonomic assignment using the sklearn classifier^32^, trained on the SILVA v138 dataset available on the QIIME2 website.

Isolate 16S Bioinformatics: The negative control had 29 sequences of one OTU and thus we subtracted 29 sequences across the entire dataset, thus effectively removing lower abundant OTUs. We consider an isolate pure if it meets the following criteria: within a single sample/isolate, 1) the most abundant sequences must make up at least 90% of an OTU; or 2) for isolates not meeting criterion 1, the most dominant OTU must be 10X more abundant than the second most abundant OTU; or 3) the most abundant OTU belongs to the same “species” as the less abundant OTUs with >95% classifier confidence. This stringent approach allowed us to detect and remove isolates that may be contaminated or simply contain artefacts of high-throughput sequencing.
